# An Open Resource: MR and light sheet microscopy stereotaxic atlas of the mouse brain

**DOI:** 10.1101/2024.03.28.587246

**Authors:** Harrison Mansour, Ryan Azrak, James J. Cook, Kathryn J. Hornburg, Yi Qi, Yuqi Tian, Robert W. Williams, Fang-Cheng Yeh, Leonard E. White, G. Allan Johnson

## Abstract

We have combined MR histology and light sheet microscopy (LSM) of five postmortem C57BL/6J mouse brains in a stereotaxic space based on micro-CT yielding a multimodal 3D atlas with the highest spatial and contrast resolution yet reported. Brains were imaged in situ with multi gradient echo (mGRE) and diffusion tensor imaging (DTI) at 15 μm resolution (∼ 2.4 million times that of clinical MRI). Scalar images derived from the average DTI and mGRE provide unprecedented contrast in 14 complementary 3D volumes, each highlighting distinct histologic features. The same tissues scanned with LSM and registered into the stereotaxic space provide 17 different molecular cell type stains. The common coordinate framework labels (CCFv3) complete the multimodal atlas. The atlas has been used to correct distortions in the Allen Brain Atlas and harmonize it with Franklin Paxinos. It provides a unique resource for stereotaxic labeling of mouse brain images from many sources.

---

We have improved the spatial and contrast resolution of magnetic resonance histology, i.e. microscopic MRI of postmortem specimens enabling quantitative assessment of strain variability in CNS structure and cytoarchitecture in the mouse^1^. We have developed a computational infrastructure that makes it practical to merge multimodal MRI data sets with full resolution whole brain, immunohistochemical volumes generated using light sheet microscopy (LSM). This has allowed us to register stained datasets highlighting many distinct classes of neurons, glia, and vascular elements into MR reference volumes that have high intrinsic geometric accuracy because every brain is imaged while still in the skull ^2, 3^. To facilitate integration within the community of investigators using mouse brain models, we translated the common coordinate framework (CCFv3) from the Allen Brain Atlas (ABA) into our refence space. In so doing it became clear that there is need for a stereotaxic atlas to correct for the inherent distortions in the ABA and many other atlases to facilitate data comparison. These distortions compromise the accuracy of estimating cell densities, structure volumes, and distances among cranial features in adult mice ^2^. A perhaps tolerable 5% inaccuracy in a linear measure, can lead to 15% volumetric error. Distortions in shape also introduce serious targeting error for stereotaxic placements. For example, an angular distortion of 5 degrees at the cortical surface, can lead to an offset of more than 500 μm at a depth of ∼6 mm. These kinds of errors become even more problematic when attempting to quantify differences in the architectures of brains of highly diverse "wildtype" strains that have brain volumes that vary from 0.32 to 0.56 cm^3^ ^4–6^. Moreover, geometrical distortions are themselves dependent on brain regions, depths from surface, and neurite and axonal orientations. This leads to striking anisotropies, and it is not possible to apply a single global scalar factor to correct data sets. This is particularly true for tissue processed using the many variants of the original CLARITY protocol ^7^. For these reasons we have produced a geometrically accurate population atlas of the brain of young adult C57BL/6J mice with spatial and contrast resolution beyond that of any previous work and have registered it into stereotaxic coordinate space using high resolution CT scans of the whole skull to ensure precise definitions of lambda- and bregma-based skull coordinates. We have adapted the ABA Common Coordinate Framework (CCFv3) that defines several hundred regions of interest (ROIs), ^8^ but we have now corrected the volumes of these regions to conform much more closely to *in vivo* volumes and coordinates.

Waxholm Space (WHS) was defined in 2010 by a working group of the International Neuroinformatics Coordinating Facility (INCF.org) as an image-based reference for coordinating mouse brain research ^9^. It provided the first isotropic 3D atlas of the C57BL/B6 mouse brain with multiple MRH volumes registered with Nissl sections and a set of 3D delineations. It was conceived as a reference space to which other atlases might be mapped enabling comparison across the many available atlases in a common space. Table S1 lists some of these existing atlases. The concept is even more relevant today and we report here the Duke Mouse Brain Atlas (DMBA) of the adult male C57BL/B6 mouse brain with significant new additions for comparative anatomy: 1. The highest spatial resolution MRH images yet acquired; 2. Assembled with a minimum deformation template using five specimens yielding the highest contrast to noise yet published; 3. With fourteen different MR imaging contrasts each highlighting unique histological and cytoarchitectural features; 4. Registered to microscopic CT images providing stereotaxic reference to cranial landmarks; 5. With high angular resolution diffusion MRI (dMRI) yielding track density images at 5 μm isotropic resolution; 6. With 180 delineations in each hemisphere consistent with the Common Coordinate Framework (CCFv3) ^1^; 7. With full resolution (1.8×1.8×4.0 μm) whole brain light sheet images from seventeen different immunohistochemical markers all registered into the same stereotaxic space; 8. With enhanced FAIR distribution through a public website designed for the very large, multidimensional image volumes.

The significance of the DMBA is four-fold: DMBA provides a geometrically accurate atlas for the C57BL/6J strain that has been widely used in molecular, cellular, and functional studies of mice. As we demonstrate here, DMBA is suitable for registration of LSM data into MRH data. It will also facilitate mapping 2D histology images from multiple sources into a stereotaxic space ^10^. The precision of alignment in the MRH and LSM is better than 50 microns and without post-hoc adjustments. DMBA can be used to estimate and compare densities of cells, volumes of brain structures, cross-sectional areas of tracts, and complexity of connectomes across the entire CNS with coefficients of error that will be limited more by image contrast and specificity of stains than by geometric errors. The combination of high spatial resolution and multimodal contrast resolution enables DMBA to harmonize many existing atlases across scales into a common stereotaxic space. The suite of contrasts derived from the mGRE and DTI facilitate label mapping in preclinical MRI studies at far higher resolution than any existing MRI atlas. DMBA immunohistochemistry at full resolution provides a path for registering intrinsic signal and cell-based atlases like the ABA and Franklin-Paxinos ^11^. DMBA enables integration of diverse stereotaxically defined physiological and optogenetic data. The range of contrasts facilitate machine driven registration across a broad range of strains beyond the C57BL/6J ^12^ at different ages in healthy and diseased animals ^3^.

## RESULTS

### Overview

The unifying core of the atlas is a library of 3D MRH volumes derived from five 90±2-day C57BL/6J male mice (**Table S2**). Multi gradient echo (mGRE) and diffusion weighted MRI (dMRI) were acquired at 15 μm isotropic spatial resolution with the brains in the skull limiting global or regional swelling or shrinkage and preventing any further volumetric distortion or tissue destruction that might otherwise accompany cranial dissection and conventional histology. The MRH volumes for all five specimens were diffeomorphically registered together increasing the contrast to noise. Fourteen different average volumes were derived from the collection of MRH scans, each highlighting different meso- and micro-scopic features of cerebral tissue (**Table S3**). Micro-CT images (@ 25 μm isotropic resolution) were registered to the MRH average volume to provide critical cranial landmarks (bregma and lambda). The brains used for the MRH volumes were then removed from the skull, cleared, labeled with fifteen different immunohistochemical markers and processed for LSM. A specimen in which cells expressing the Thy-1 cell surface antigen were visualized with genetically encoded yellow fluorescent protein (YFP) ^13^ was also scanned with the MRH protocol but was not included in the MRH atlas. The specimens summarized in **Table S4** were scanned using structured plane illumination microscopy (SPIM) at cellular resolution (1.8×1.8×4.0 μm). These light sheet microscopy (LSM) images were registered at full resolution to the MRH volumes correcting the geometric distortion common in LSM and placing them in the stereotaxic space ^2^. The registered dMRI volumes in the common space were used to create super resolution (5 μm) track density images ^14^. A subset of isotropic 3-dimensional labels from the common coordinate refence framework (CCFv3) ^8^ was registered to the average MRH volumes placing these widely used labels in the stereotaxic space.

### Average MRH atlas

In previous work we have described dMRI and mGRE images acquired at 15 μm isotropic resolution ^3^. This new stereotaxic atlas expands on that in four critical ways by creating average dMRI and mGRE volumes with much higher contrast to noise, mapping everything into stereotaxic space with cranial landmarks from micro-CT, generation of an average light sheet atlas (NeuN) in the same stereotaxic space, and the addition of 17 LSM volumes (in the stereotaxic space) with unique IHC stains. Figure 2 demonstrates how the contrast resolution has improved in the MRH.

**Figure 1.**
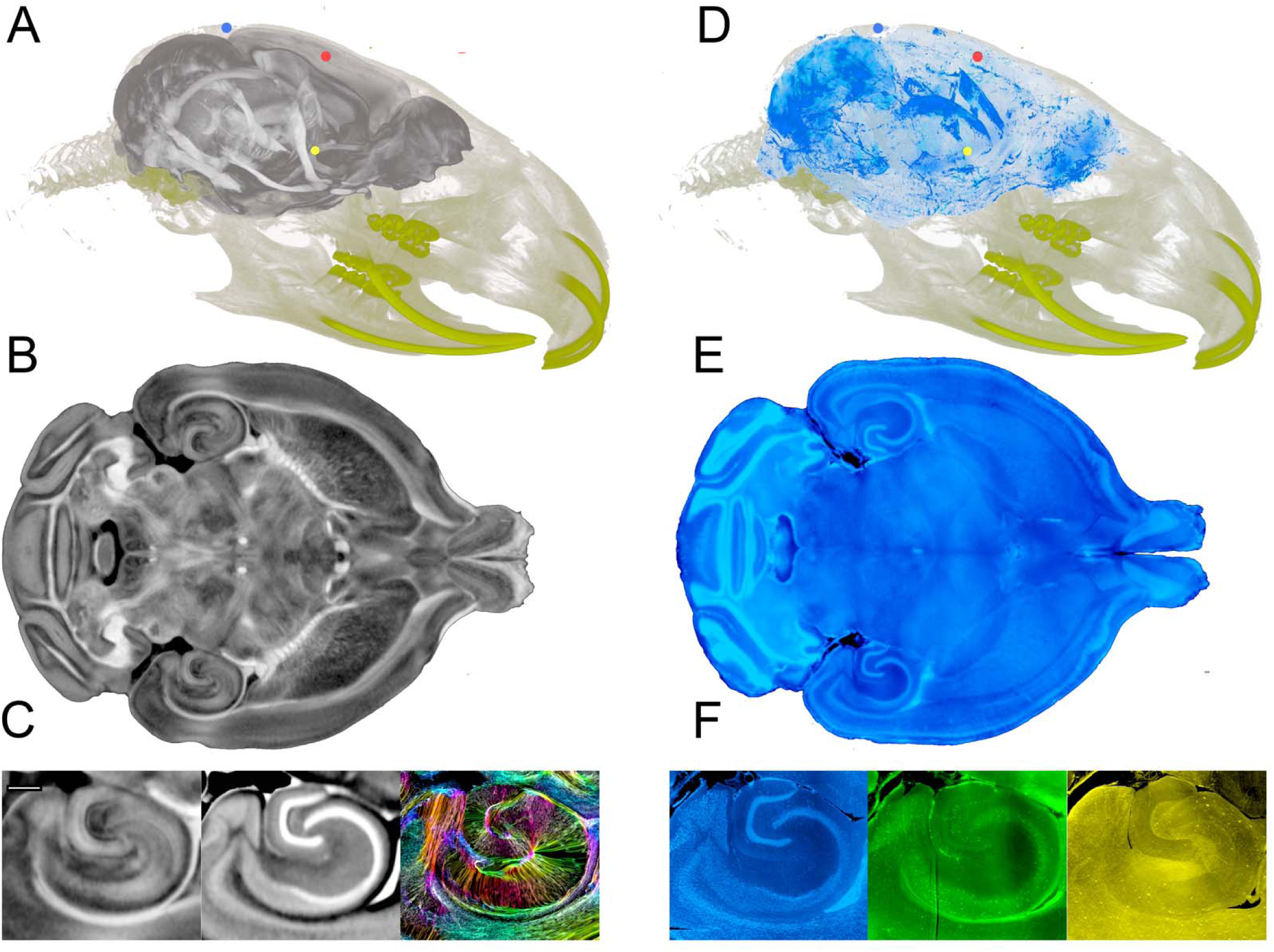
The Duke Mouse Brain Atlas (DMBA), a multimodal stereotaxic atlas of the mouse brain. DMBA is a collection of 31, 3D MR histology and Light Sheet Microscopy volumes of the adult male C57BL/6J mouse brain registered to a common stereotaxic space consistent with Franklin and Paxinos ^11^. The stereotaxic space (**A–C**) is defined by postmortem MRH scans of five males (90±2 days) using extended acquisition (132 hours/specimen) at 15 µm isotropic resolution. Scans were acquired with brains in the skull minimizing geometric distortion. Multigradient echo and DTI sequences were used generating 14 different volumes, each highlighting different meso- and micro-scopic features of cerebral tissue. The five specimens were registered to a minimum deformation template to create a collection of average 3D volumes with exceptional spatial and contrast resolution. **A**) Volume rendered Micro CT images of the skulls were registered to the 3D average MRH Fractional Anisotropy (FA) volumes providing cranial landmarks (lambda and bregma; blue and red dots, respectively; the anterior commissure is marked with a yellow dot). **B**) Representative 15 μm axial slice from the FA volume demonstrates in high definition the myeloarchitecture of the brain. **C**) Comparisons of the hippocampal region defined in 3 of the 14 different MRH modalities: **left:** fractional anisotropy (FA); **middle:** mean diffusivity (MD); **right:** track density images at 5 μm isotropic resolution showing fiber density and orientation. Eleven additional 3D volumes derived from the MRH acquisitions provide different histological detail (**Supplement Table S3, Figure S1**). **D**) A complementary collection of light sheet microscopy images was generated by clearing brains after MRH scans, staining with a selection of immunohistochemistry markers, and scanning using selective plane illumination microscopy. These volumes were corrected for distortion arising from dissection and clearing by affine and diffeomorphic registration to the MRH volumes (in the skull). **D)** shows the geometrically corrected whole brain light sheet volume for NeuN mapped into the 3D CT volume. **E)** is a representative 4 μm axial slice (1.8 μm in-plane resolution) from the same NeuN volume at full resolution (∼300 GB/volume). Closer inspection of this full resolution NeuN image and two additional stains—from a total of 17—shows the same hippocampal region as in **C** but with spatial resolution sufficient to resolve single cells. **left:** individual cells positive for NeuN (RBFOX3 protein); **middle:** cells positive for parvalbumin (PV); **right**: cells positive for neuropeptide Y (NPY). The scale bar in C is 0.4 mm.

**Figure 2.**
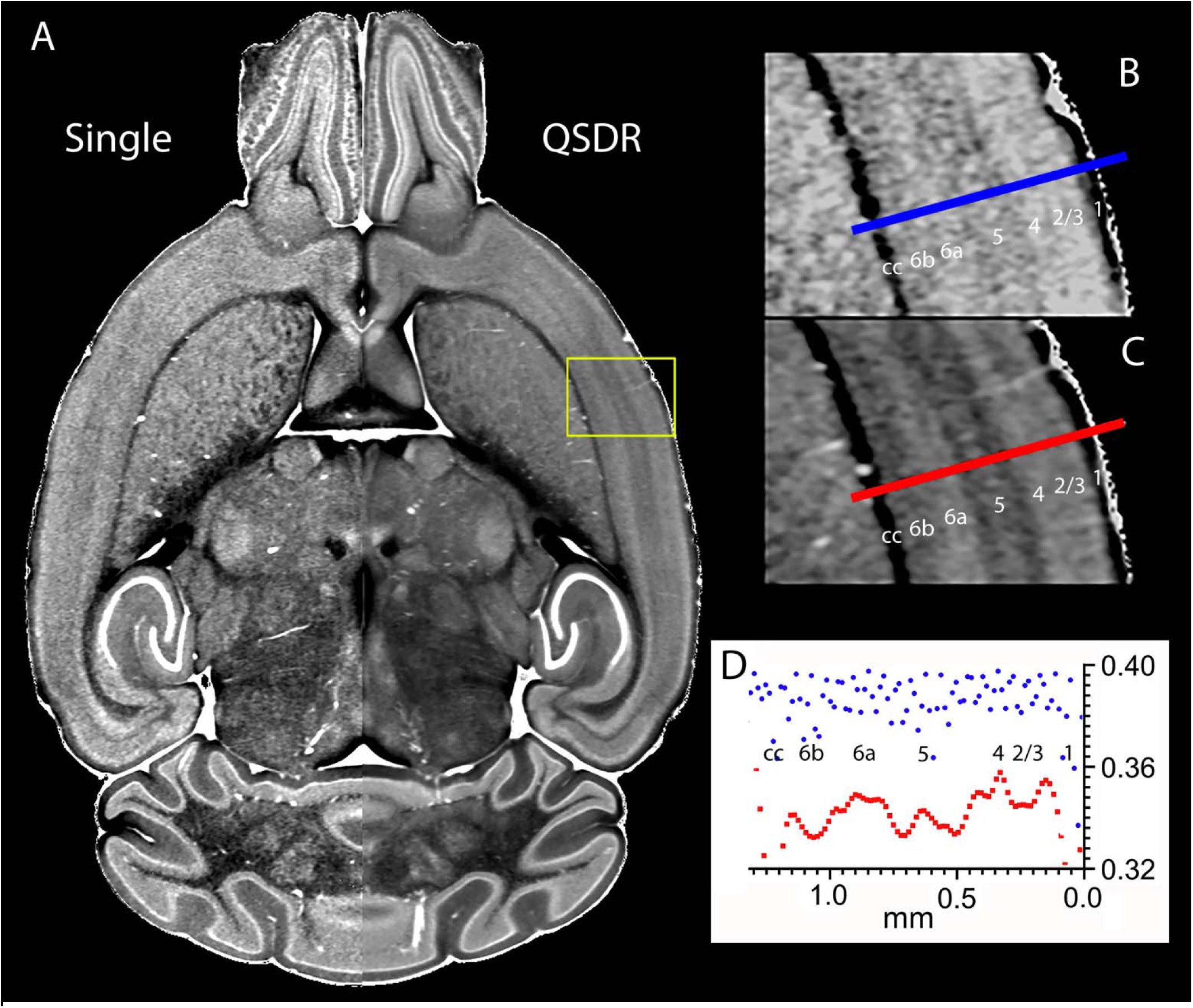
Precision registration improves image contrast and signal to noise. A) A single animal (left side, case200302-1:1, MDT contrast) compared to the average of five cases (right side; QSDR). B) Neocortex (primary somatic sensory cortex) highlighted in A for the single specimen, and C) for the same region from the MDT average image, with traces defining a plot across both and numerical labels to indicate cortical layers (cc; white matter of corpus callosum). D) The gray scale value across the single specimen (blue) is noisy, with cortical layers poorly differentiated. The plot across the same area in the MDT average (red) clearly defines cytoarchitectonic borders between cortical layers and even sublamination in cortical layer 5. Similarly, other cytoarchitectural features of thalamic nuclei and cerebellar cortex, including the Purkinje cell layer, are more distinct in the average image.

The improvement in contrast to noise is similar across all the average images resulting in one of the most unique features of this atlas, the definition of meso-scale structures seen in **Figure 3**. Hippocampal and cerebellar layers not seen in the gradient recalled image (**Figure 3A**) become readily visible in the diffusion weighted image (DWI), **Figure 3B**. The Purkinje cell layer in cerebellar cortex is visible in the mean diffusivity (MD) image (**Figure 3D**) and subtle features of cerebellar white matter in the lobules is revealed in the fractional anisotropy (FA) image (**Figure 3G**). The color FA image in **Figure 3H** provides evidence of specific orientation of these fascicles. **Table S3** summarizes the fourteen volumes included in the MRH atlas. Most of the volumes are provided with the skull stripped as this facilitates their use in automated registration to other atlases. An average mGRE without masking is provided to help mapping *in vivo* MRI data. **Figure S1** shows a full field representative axial plane at full resolution allowing the reader to examine a larger region of interest interactively.

**Figure 3.**
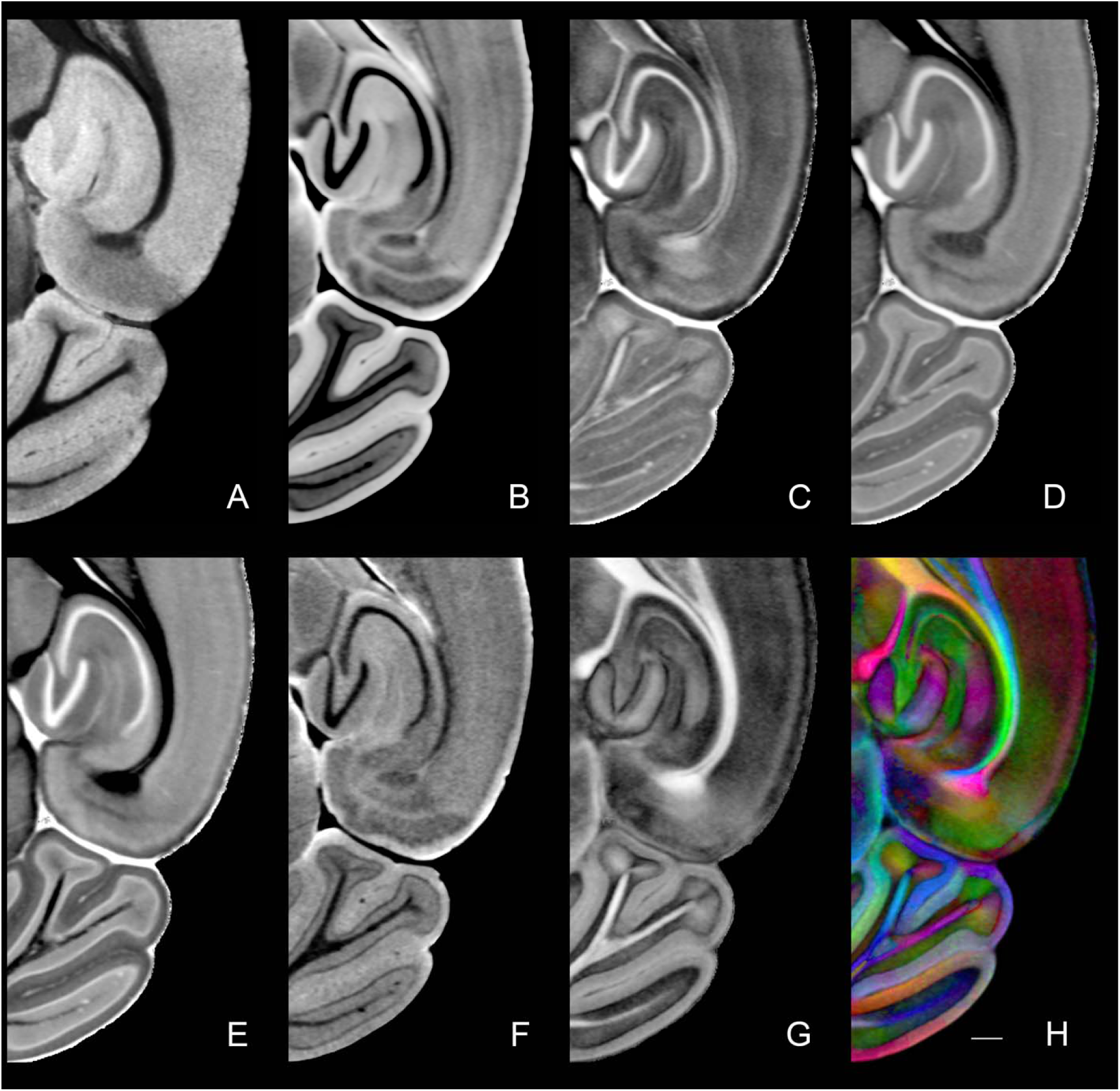
Magnified sections from eight of the volumes from the average MRH atlas show the marked improvements in definition of cytoarchitectural boundaries. **A**) Average of four echoes from the mGRE; **B**) Diffusion weighted image (DWI); **C**) Axial diffusivity (AD); **D**) Mean diffusivity (MD); **E**) Radial diffusivity (RD); **F**) ISO from GQI algorithm; **G**) Fractional anisotropy from DTI algorithm; **H**) Color FA from DTI algorithm. The scale bar in H is 0.4 mm.

Averaging five specimens produces a few confounds. High contrast penetrating vessels in similar but not identical areas will "shine through" in the average atlas giving one an impression of higher vascular density, as shown in **Figure S2**. **Figure S3** demonstrates the blurring one might expect by focusing on modular features of brain cytoarchitecture, such as the glomeruli in the olfactory bulb. They are discrete and well defined in the images from the individual specimens before averaging. After averaging, they are blurred; but the layers of the olfactory bulb defined by intrinsic cytoarchitectural features are enhanced.

### Stereotaxic CT

Five 3D micro-CT images were acquired at 25 μm isotropic spatial resolution to provide critical cranial landmarks (bregma and lambda) for defining stereotaxic space. Each CT volume was aligned to the average MRH volume using a full affine transform to estimate the potential biological variability one might encounter in a real-world stereotaxic environment (See methods). The most significant outlier was a displacement of 450 μm between a single volume and the average of the five. The DMBA includes two collections (MRH and LSM) of data in the common space (**Figure 1**): A volume rendered view of the CT data combined with the DWI (**Figure S4A**) includes the incisors used in stereotaxic frames to establish the coronal plane. **Figure S4B** provides a midsagittal slice of the combined CT and DWI focused on the DWI with bregma marked with a red dot and lambda with a blue dot. A third landmark in the center of the anterior commissure (AC, yellow dot) defines the coronal plane as the plane intersecting bregma and the AC. The coronal plane for lambda is shown in **Figure S4C** and the coronal plane for bregma is shown in **Figure S4D**. We have chosen this orientation to be consistent with that used by Franklin and Paxinos ^15^. **Figure S5** shows the midsagittal plane for the DMBA DWI and the sagittal Nissl image from ref. 15 (Plate 104). The estimate of the "flat skull" angle has been marked for both along with the angle of the corpus callosum, which differs by less than 0.5 degree between atlases. **Figure S6**, a comparison of coronal planes through bregma for ref. 14 and the DMBA confirms the orientation of the two atlases is very close. The two images in **Figure S6** have been scaled to allow visual comparison of anatomy but they are not scaled to correct for the shrinkage that accompanies the preparation of histological sections and the treatment of those sections to produce a Nissl stain. Distortion in the Nissl is evident in the aspect ratio of the two images. The height/width is 1.31 (ref. 15) to 1.21 (DMBA) which is consistent with the differential change in height/width that usually accompanies removal of the brain from the cranial vault. **Figure S5** and **S6** confirm that the orientation of the DMBA is very close to that of ref. 15.

### Labels

We have adopted a version of the common coordinate framework (CCFv3) ^8^ as the default labels and ontology for the atlas. CCFv3 delineates a total of 461 structures. The label set included here starts with this label set and ontology. Our use of the atlas employs pipelines for labeling MRH volumes of many strains other than the C57BL/6J upon which CCFv3 is based and translating labels between MRH and LSM volumes with widely different histochemical stains ^2^. Smaller, poorly registered regions of interest (ROI), such as neocortical laminae or subdivisions of subcortical nuclei, pose a technical source of noise that can impact the validity of the registration ^1, 3^. We describe an intermediate path with the reduced CCFv3 label set (RCCFv3) in which related structures with volumes < 0.1 mm^3^ have been aggregated into larger volumes of which the structures are subcomponents. **Figure 4** shows the RCCFv3 label set on three planes of two of the MRH volumes highlighting the fact that the different contrasts derived from the dMRI volumes (MD and FA) provide complimentary information that can assist the registration algorithms. The full list of structures, abbreviations, and relations between CCFvs and RCCFv3 can be downloaded with the atlas. The accompanying JSON file (RCCFv3_JSON) provides explicit description for all the columns.

**Figure 4.**
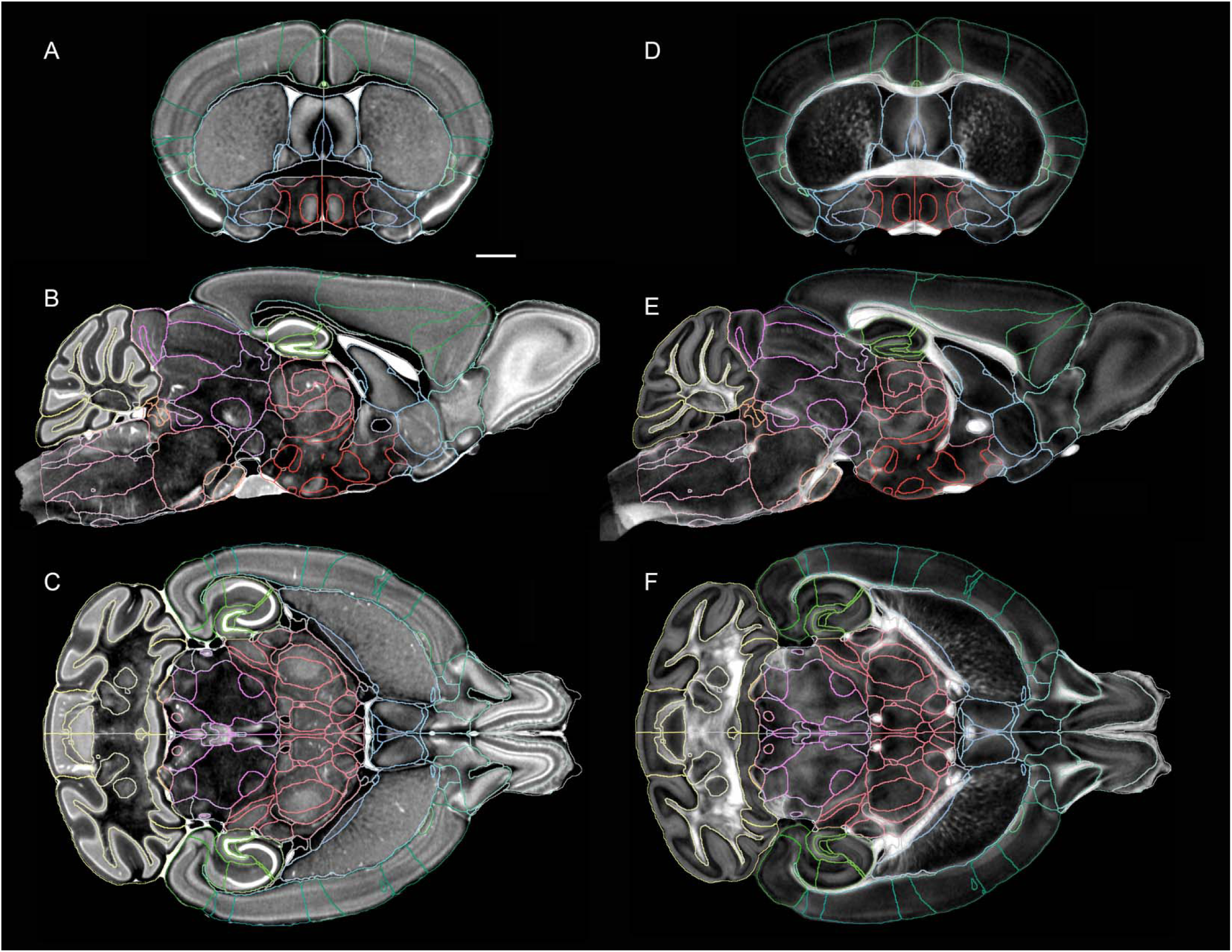
Multiple dMRI volumes provide complimentary cytoarchitectural support for labels. Previously ^1^, we found that our workflow could reliably register ROI with volumes greater than 0.1 mm^3^ across four strains (C57BL/6J, DBA/2J, CAST/EiJ, and BTBR T<+>Itpr3 <TF>/J) with coefficients of variation between left and right hemisphere of less than 5%. The relations between structures and related subvolumes in RCCFv3 and the similar structures in CCFv3 are included in the Excel data sheet available with the download of the atlas. There are 180 unique ROI in each hemisphere for RCCFv3. The RCCFv3 labels are shown on the MD volume (**A–C**) and the FA volume (**D–F**). The labels are isotropic, and color coded to correspond with the standard used in the ABA (https://atlas.brain-map.org/atlas). The scale bar is 1 mm.

### Diffusion Tensor Data

The scalar images of anatomy described above are derived using the DTI algorithm and analogous GQI algorithm ^16, 17^ (See **Table S3**). These algorithms provide the basis for defining whole brain connectivity ^18–21^. Several quantitative anatomic features can be derived from the connectome. We have used the GQI algorithm to derive these metrics since it addresses the complication of crossing and/or touching fibers in a voxel through the use of high angular resolution diffusion imaging ^22^. There is a robust literature addressing tradeoffs in spatial and angular resolution ^23^ ^24, 25^ ^26^ and the efficacy ^27^. The connectome data in this atlas addresses the enormous differences between clinical scans acquired at 2 mm resolution (8 mm^3^ voxels) over a 1.5 kg human brain and 15 μm^3^ voxels that are 2.4 x 10^6^ times smaller over a 435 mg mouse brain. The resolution index (angular resolution/voxel volume) ^28^ in the human connectome is 527 angular samples / mm^3^. In our previous work we found good agreement between DTI and the results of conventional *in vivo* tract-tracing experiments using retroviral tracers in a mouse with DTI datasets at a resolution index of 1.51 x 10^6^ angles/mm^3^ ^21^. The resolution index for the DMBA at 3.2 x 10^7^ samples / mm^3^ is more than 20 times that of our previous work making this the highest resolution connectome of the mouse brain available to date. **Figure S7** shows the connectome of one of the individual specimens and that of five specimens that have been averaged together using the QSDR method ^17^. Improvement in the sensitivity is clear, especially for connections within and across the cerebral hemispheres (intracortical and subcortical fiber tracts, including commissural tracts). Such connectome data are representative of ongoing work to validate the node-/edge-specific results in quantitative terms and in relationship to experiments with axonal tracers—a necessary step toward creation of a white matter atlas of the mouse brain.

### Population NeuN

Contrast defining histological features, such cytoarchitectural boundaries and myeloarchitectural structures, in MRH and LSM images is dependent on completely different phenomena (e.g., diffusion of water vs. immunohistochemical stains). This presents a challenge to algorithms matching similarity or strong mutual information. We have optimized the registration between MRH and LSM based on DWI and MD and NeuN ^2^. **Figure S8A** shows a comparison of the DMBA MD image with a minimum deformation template created from 4 LSM NeuN images (**Table S4** Specimens 220905_1-4). While there is not a precise correspondence between the signal in the two images, there are several regions well distributed throughout the volume with consistent signal relations that help drive the registration. For example, cell-dense regions yield high signal in MD. As a marker of neuronal nuclei, one should expect brain structures with high neuronal density should correspond to those high-signal regions in MD images. This is indeed evident across the brain with striking examples in the granule and pyramidal cell layers of the hippocampus, the granule cell layer of the cerebellar cortex, the granule cell layer of the olfactory bulb, and the granular and supragranular layers of the neocortex. **Figure S8 B, C** highlights the alignment of the 15 μm MDT NeuN (B) with an individual volume (220905-1_2) at full resolution (1.8×1.8 μm), where individual neuronal nuclei are clearly seen (C).

### Grids

Six different grids are included with the atlas (See Methods) all indexed to bregma. Three grids in xy, xz, and xy planes are defined at 1 mm spacing. The xy grid is visible in **Figures 5** and **S8A** with the coronal and sagittal planes passing through bregma highlighted in thicker lines; other gridlines are equally spaced at 1 mm intervals off these planes. A second collection of 3 grids with 200 μm spacing is included, again indexed to bregma (see **Figure S8B and C**).

**Figure 5.**
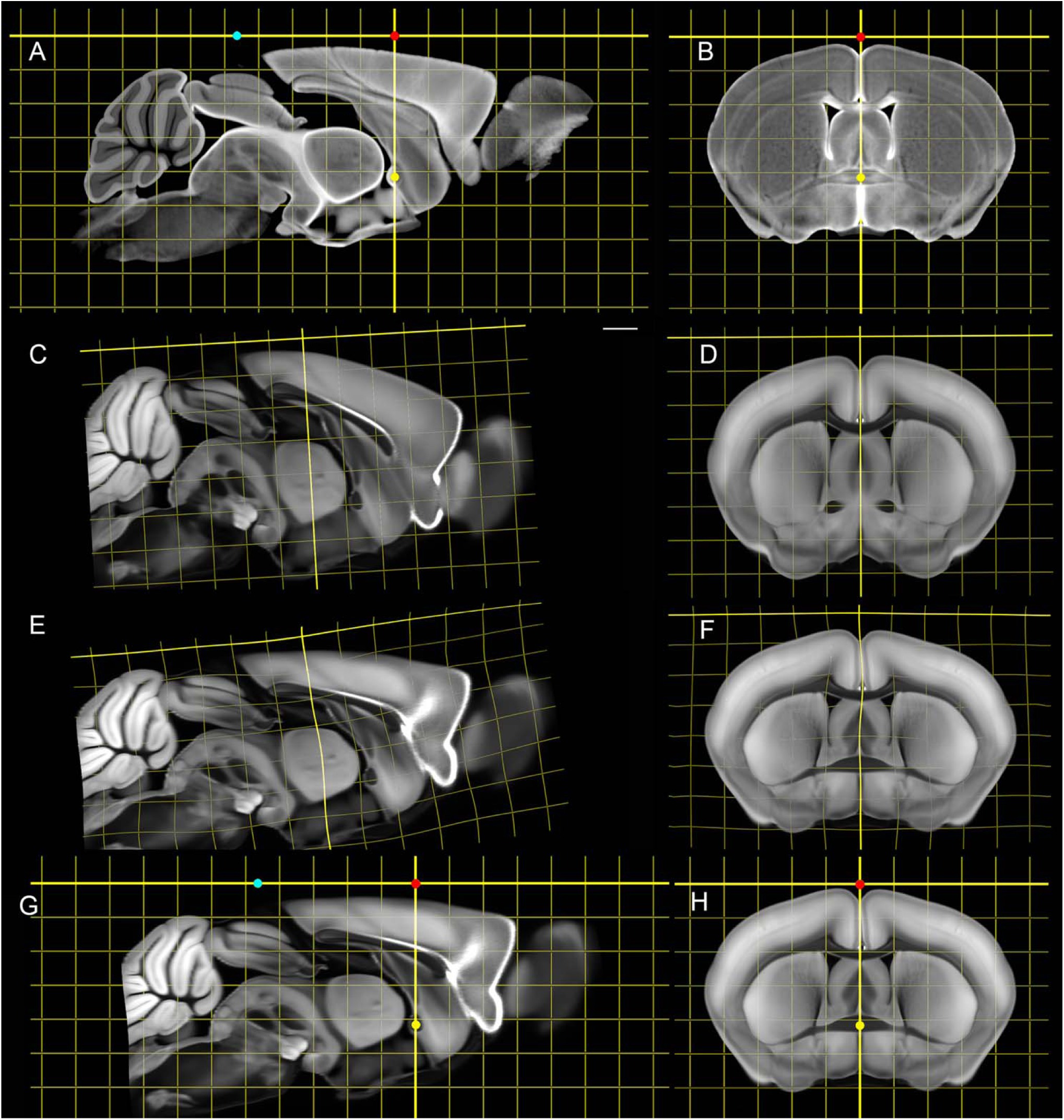
Mapping the ABA into the DMBA corrects the geometric distortion and places the ABA in stereotaxic space. The ABA auto fluorescent volume was down sampled to 15 μm and registered to the DMBA-DWI volume using both affine and diffeomorphic transforms (see Methods). **A**) 3D regular grid was imposed into the DMBA-DWI space in both **A** and **B**. Since bregma and lambda are not defined in the ABA, the first step was to estimate their locations in ABA space. An affine and diffeomorphic transform was applied to the ABA volume to map it into DMBA. The transforms were inverted and applied to the DMBA skull landmark set, mapping bregma into ABA space. A regularly spaced grid was then created in ABA space grid with an origin at bregma. **C** and **D** shows the ABA, and the constructed ABA space grid, after affine warping to DMBA, demonstrating that there is global shrinkage in the ABA and alignment error of the z axis relative to DMBA. It is not surprising that this estimate returns a coronal slice at bregma in which the plane misses the midsagittal crossing of the anterior commissure (**D**). **E** and **F** shows the impact of the affine and diffeomorphic transform highlighting the fact that the geometric corrections to bring ABA into DMBA space are not uniform. The ABA is assembled from a series of slices from multiple brains so this may be due to shrinkage of the tissue or an artefact of the reconstruction. The z axis is tilted at the midpoint by ∼4 degrees. This is consistent with behavior we have seen when the brain is removed from the cranial vault with more significant distortion in olfactory bulbs and the brainstem ^2^. The corrected volume is stretched along z axis by ∼4%. **D** and **F** highlight differential swelling in the coronal plane. The swelling is 16.3% along the y axis and 5.7% along the x axis. **G** and **H** show the final result with the ABA now in the same stereotaxic frame as DMBA, with corrected geometry and orientation, and external landmarks for bregma and lambda. The results are consistent with similar work of Perens et al^29^. The scale bar is 1 mm.

### FAIR Distribution

The data reported here is available at CIVMImageSpace (https://civmimagespace.civm.duhs.duke.edu/). “© 2024 Duke University. All Rights Reserved. The data was produced at the Duke School of Medicine, Department of Radiology and Pratt School of Engineering, Department of Biomedical Engineering and is being made available under a Creative Commons CC BY-NC-SA 4.0 License. See this URL for additional information <u>CC BY-NC-SA 4.0</u> <u>Deed | Attribution-NonCommercial-ShareAlike 4.0 International | Creative Commons</u>” CIVMImageSpace has been constructed to handle very large multilayered (4D) image arrays and provide users the option of review to determine which data they would like to download. The site supports users who want full resolution and those who will benefit from lower resolution, more compact data. The aggregate library is ∼13 TB. All data is displayed in RAS orientation. The data is grouped with FAIR standards and FAIR compliant metadata and data files. Images are available to download as NRD files. N5 files are displayed on the web site using a hierarchical display that allows users a more seamless interaction when there are bandwidth limitations. A web enabled viewer (Neuroglancer) supports interactive review *prior to download.* This facilitates exploration without waiting for large volumes to download. The user chooses desired data and launches automated large queues. Consistent image headers across all volumes for RAS orientation facilitate interoperability of the volumes across multiple software packages.

### Applications

The DMBA has been envisioned as a resource to bridge the gap across many neuroscience communities, including (1) preclinical imaging (MRI) studying mouse models of human disease; (2) neuroscientists requiring precise and accurate coordinates for stereotaxic localization in experimental protocols; (3) investigators needing to correct geometric distortion in light sheet and confocal microscopy; (4) those using multiple 3D volumes and needing to integrate data from diverse sources into a common reference space. We demonstrate the utility of DMBA in two such examples.

### The Allen Brain Atlas (ABA)

The Allen brain atlas (ABA) ^8^ has become a unifying reference and resource across many domains of neuroscience. One major limitation in the ABA is forthrightly stated: " … at present, CCFv3 is not suitable for determining stereotaxic coordinates." ^8^. We demonstrate in **Figure 5** the use of DMBA in making multiple improvements to the ABA. Thus, DMBA corrects geometric distortion in the ABA while provide cranial stereotaxic landmarks and additional sources of contrast to isolate mesoscale brain structures. Importantly, the DMBA makes the internal structure of mouse brain more accessible to preclinical MRI users, and it makes the multidimensional properties of MRH accessible to neuroscientists who are not yet familiar with MRI-based applications in mouse neuroscience.

Table S4 in Wang et al. ^8^ provides a summary of the average volumes of the structures in the ABA. The authors note that the atlas is not stereotaxic, so the absolute volumes are not accurate. The transforms between ABA (**Figures 5C, D**) and the stereotaxic correction demonstrated (**Figure 5G,H**) were applied to the ABA labels to estimate the changes upon stereotaxic correction. **Figure S9** compares nine of the larger volume changes in that transformation showing volumes in ABA space are up to 8.5% larger and there is a relatively large variation in the change. The differential swelling along x and y and shrinkage along z lead mean that comparison of relative volumes in ABA space is problematic. **Figure S10** demonstrates that there are both decreases and increases in ROI volume upon transformation to stereotaxic space and the correction ranges from ∼ -25% to +15%.

The correction for differential shrinkage in z and swelling in x and y in the ABA places the ABA and FP atlases in close alignment due to the angular correction along z, a distortion that is not as prominent in FP (See **Figures S5 and S6**). But now the agreement between coronal sections in the corrected ABA and FP is very good. Perhaps more importantly, the correction improves estimates of structural volumes crucial in following quantitative volume change in brain development, age-associated neurodegeneration, and across strains.

A final benefit to the correction of the ABA arises from the additional new information from MRH to assist imaging pipelines in delineating mesoscale anatomy, as depicted in **Figures S11 and S12** There are several existing pipelines that map the ABA (and labels) to optical images (confocal and LSM) ^2, 29–32^. These work well for the healthy male C57BL/6J. The challenge comes with different strains and ages. A catalogue of MRH and LSM volumes in the common space with labels provides *multiple* templates for anatomic delineation where boundaries are less clearly defined. In **Figure S11** cortical layers are better defined in the DMBA-MD, AD, and DWI volumes than the ABA. Boundaries of thalamic nuclei are much better circumscribed in the DMBA-MD with complimentary information from the DWI than the same regions in the ABA. The fiber orientation displayed in the color FA (**Figure S11G**) resolves fimbria (yellow) from stria terminalis (blue) and internal capsule (red) in a region devoid of signal in the ABA atlas. In the magnified hippocampus in **Figure S12** differential contrast in DMBA-AD, DWI and FA help delineate and differentiate the cell-dense layers (granular layer of the dentate gyrus and pyramidal cells layers of the CA fields) from the neuropil-rich layers of the hippocampal divisions. Moreover, these DMBA MRH contrasts reveal sublaminar structure in the CA fields, such as the differentiation of the stratum lacunosum-moleculare from the stratum radiatum in CA1 (see **Figure S12 F** and **G**).

**Figure S13** highlights the new anatomic insight provided by the track density images. Calamante et al reported the super resolution method in 2010 ^14^. They compared super resolution TDI in a mouse brain to conventional histological sections stained for myelin using Gallyas silver stain in a mouse brain in 2012^33^. Their data was acquired with a Stesjkal Tanner spin echo sequence encoding at 100 um isotropic resolution with 30 directions at b=5000 s/mm^2^. Super resolution construction allowed them to extend the spatial resolution down to 20 μm. The data in the DMBA was acquired with 15 μm^3^ voxels that are ∼ 300 times smaller with 108 angular samples (resolution index that is more than 1000X larger) allowing us to use a step size of 5 microns. The light sheet volume (Specimen 200316-1) stained with myelin basic protein was registered using the NeuN image, which was one channel of the acquisition (**Table S4**). The data in **Figure S13** employs the method described by Calamante to address the dynamic range by constraining the length of the local neighborhood contributing to the display to <1 mm.

Calamante et al. ^33^ points out the challenge in the comparisons between TDI and myelin arising from the inherently different mechanisms giving rise to each image. Structures other than myelinated axonal fascicles can readily yield signal in the TDI, as discussed above for neocortex, Similarly, a comparison of Thy-1 expression in the hippocampus with TDI signal indicates that TDI is capturing anisotropy associated with the organization of neuronal dendrites and axonal projections (**Figure S14**).

### Multiple light sheet volumes mapped to the same steoreotaxic space

The stereotaxic atlas allows one to correct for differential processing distortions supporting comparison of true cell density across multiple specimens^2, 34^. To demonstrate this potential, four specimens (Specimens 220905-1, 2, 3, and 4) were processed with immunohistochemical labels summarized in **Table S4**. They were mapped to the DMBA space through the NeuN MDT (See Methods). The precision of alignment is demonstrated in **Figure 6**. Since the 4 volumes have been corrected for geometric distortions from extraction from the skull, tissue clearing and differential staining, it is possible to make quantitative comparisons of *neuron density* using the methods outlined in ^34^. **Figure S15** Show the corrected neuron density in CA1, CA3, Subicullum, and LG for the all the specimens. The spread in the density derived from the modified stereological fractionator provides insight into the heterogeneity of the neurons in each ROI.

**Figure 6.**
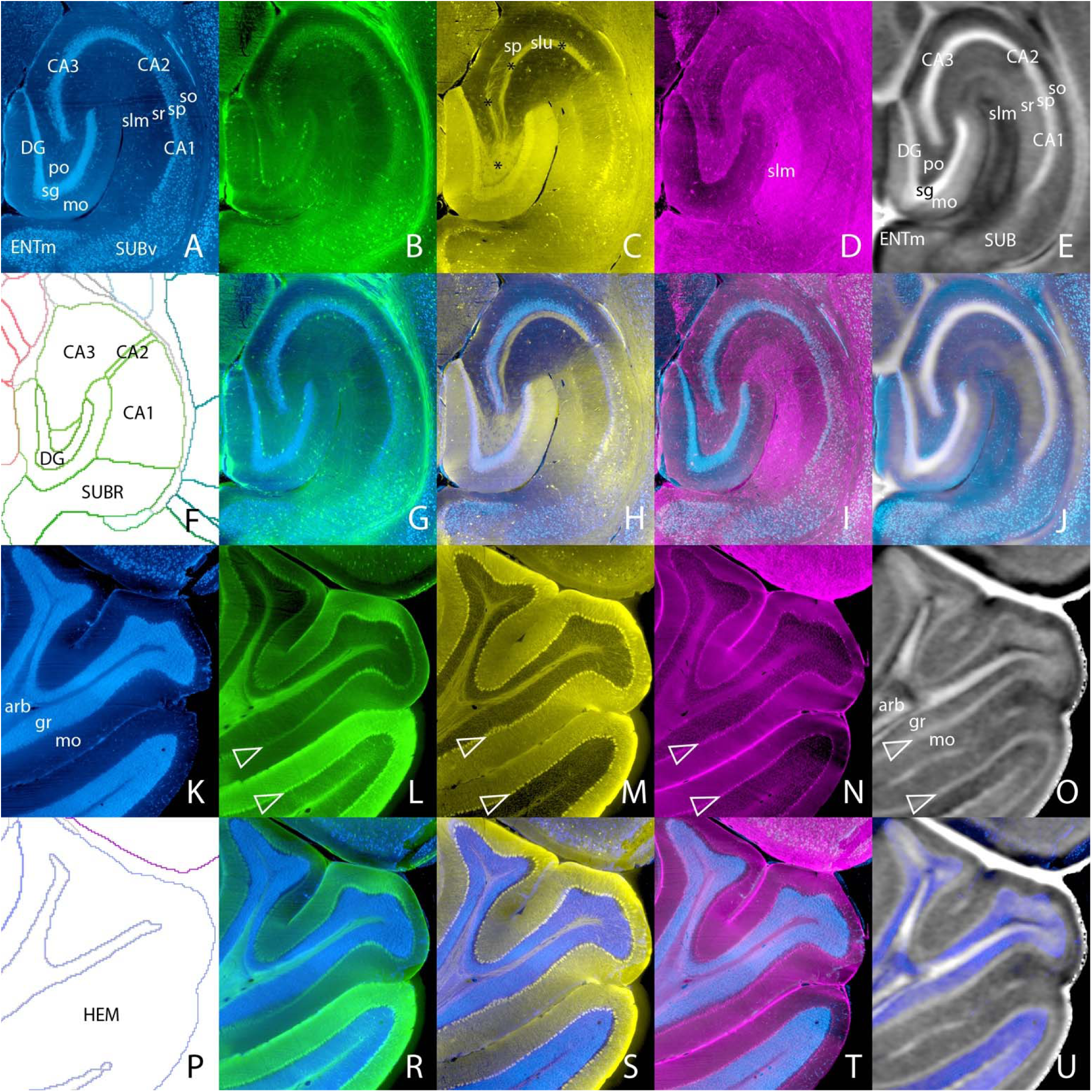
Representative light sheet images from different animals co-registered in stereotaxic space with magnetic resonance histology (MRH). Images are in the axial plane from Z = -3006 mm; same as in Figure S8 where grid lines indicate scale; light sheet images are 4 µm thick and the MRH images are 15 µm thick. The top two rows show the hippocampus, and the bottom two rows show lateral cerebellar cortex (simple and ansiform lobules). Panels (A) and (K) show NeuN expression, with Allen Brain Atlas labels; (B) and (L), parvalbumin; (C) and (M), calbindin; (D) and (N), neuropeptide Y; (E) and (O), axial diffusivity MRH; (F) and (P) show DMBS atlas labels. In the second and fourth rows, the NeuN volumes are displayed with the other light sheet and axial diffusivity volumes. In the second row, note the registration within 20 µm of the cell-dense layers of the hippocampus across volumes. In (H), note the location of the pyramidal cell layer of CA3 within the distribution of mossy fibers of dentate granule cells [marked by asterisks in (C), including the slu], which are immunoreactive for calbindin. In the cerebellar cortex, note the registration of the granule cell layer and the Purkinje cell layer (marked with open triangles) across volumes, including the axial diffusivity volume where the Purkinje cell layer gives rise to a distinct band of moderate signal. Abbreviations: arb, arbor vitae; CA1-3, Ammon’s horn fields 1-3; DG, dentate gyrus; ENTm, entorhinal cortex medial part; gr, cerebellum granular layer; HEM, cerebellum hemisphere; mo, cerebellum molecular layer; mo, dentate gyrus molecular layer; po, dentate gyrus polymorph layer; sg, dentate gyrus granule cell layer; slu, stratum lucidum; slm, stratum lacunosum-moleculare; so, stratum oriens; sp, pyramidal layer; sr, stratum radiatum; SUBR, subiculum region; SUBv, subiculum ventral part.

## Discussion

Neuroscience spans scales from live animal studies (e.g., electrophysiologists and preclinical MRI @ 0.1 mm^3^ to the cellular level e.g The Brain Initiative Cell Census Network (BICCN) at 1 μm^3^ ^35–38^. Regional volume estimates are crucial for preclinical MRI and microPET studies of aging and neurodegeneration. Geneticists need atlases that will translate across strains^1^. There have been numerous creative approaches to address this cross disciplinary need ^29, 32, 39–45^. The DMBA assembles in a single stereotaxic volume the best of all these previous efforts with MR histology with spatial resolution at least an order of magnitude higher than any previous work, with the highest contrast resolution in 14 widely varied sources of anatomic contrast, with diffusion tensor data with a resolution index more than 2 orders of magnitude beyond anything published previously, with full resolution light sheet volumes of 17 different molecular and cellular stains and a subset of the CCFv3 delineations. The breadth and depth of the DMBA can be used to bring other atlases into the same stereotaxic space to integrate differing ontologies and make more accurate and biologically meaningful comparisons.

Expansion of the DMBA is under way. The raw file from which connectomes can be generated has been registered to the ABA retroviral tracer volume to generate a white matter atlas. MRH volumes have been acquired on at least 30 different strains of mice at different ages. Additional IHC volumes have been acquired and registered to the common space. More than 300 specimens have now been mapped into the stereotaxic space with RCCFv3 labels in DMBA. Training with over 100 specimens has begun to support automated AI segmentation. We are confident that DMBA can become a critical tool in integrating mouse brain studies across the scales that span the ever-increasing breadth of contemporary neuroscience.

## Methods

### Specimen Preparation

All animal procedures were approved by the Duke Institutional Animal Care and Use Committee. Male C57BL/6J mice and adult B6.Cg-Tg (Thy1-YFG/HJrs/J ) were purchased from Jackson Laboratories. All animals were perfused with 10% Prohance (Gadoteridol) in buffered formalin. Prohance is a chelated gadolinium compound commonly used in clinical MRI as a contrast agent. It is used as an active stain in MRH to reduce the spin-lattice relaxation time (T1) from 1800 to ∼100 ms ^46, 47^. Animals were anesthetized to a surgical plane with pentobarbital. A 21-gauge needle connected to a peristaltic pump was inserted in the left ventricle. Blood was flushed using a 0.1% heparin saline solution, followed by perfusion with the Prohance/formalin mixture for ∼ 6 min. Heads were placed in buffered formalin for an additional 24 h. Mandibles were removed, and skin and muscle removed to allow use of a smaller radiofrequency coil, and brains in the cranium were placed in an 0.5% Prohance/buffer solution and allowed to equilibrate for at least 3 weeks ^48^.

### MR Acquisition

MRH images were acquired on a 9.4T/89 mm vertical bore magnet with an Agilent Direct Drive console (Vnmrj 4.0). A Resonance Research (Billerica, MA) Model BRG-88_41 coil provided peak gradients up to 2500 mT/m. The brain was placed in a solenoid coil constructed from a single sheet of silver foil. Two imaging sequences were used; 1) a multigradient echo (n = 4) sequence (TR/TE = 100/4.4 ms) and a Stesjkal/Tanner spin echo sequence ^49^ for diffusion tensor imaging (DTI) (TR/TE = 100/12-19 ms). Both sequences employed phase encoding along the short axes of the specimen (x and y) with the readout gradient applied along the long axis of the specimen (z). The DTI sequence used b values of 3000 s/mm^2^. Both sequences were accelerated using compressed sensing with an acceleration factor or 8X ^50^. A total of 108 different volumes were acquired with the b vectors of the volumes spaced equidistant on the unit sphere. A baseline (b_0_) image was acquired after each ten diffusion encoded volumes. The baseline volumes were averaged together creating a template to which all the diffusion weighted images were aligned. Alignment was accomplished using ANTs ^28^ ^51^.

### DMBA MDT Creation

The DMBA atlas is an average of five C57BL/6J male mice (90±2 day) created by our SAMBA pipeline ^52^ which is based on the Advanced Normalization Tool set (ANTs) ^51^. All registration was performed using the DWI contrast. Volumes were first masked to remove the skull. Registration was initialized by a rigid alignment to our existing reference atlas ^3^. An initial MDT template was created using a pairwise affine registration. For each volume, an affine transform was calculated to every other volume in the group. These were averaged together using the ANTs averageAffineTransforms.exe command to create an initial MDT whose volume is the average of all the individuals. We then started an iterative diffeomorphic registration process where, in each iteration, all specimens were diffeomorphically registered to the previous template using the Cross Correlation metric and Symmetric Normalization (SyN) registration. The previously computed rigid and pairwise affine transforms were used as initializers. We applied these transforms and averaged the resulting volumes together to create the next template. This was repeated six times. Once we generated the final template, we cropped the volume and rigidly warped it to a stereotactic space, based on skull landmarks derived from the micro-CT scan.

### CT Acquisition

CT data were acquired on a Nikon XT H 225 ST high resolution CT scanner in the Duke shared material instrumentation facility (SMIF). Specimens were mounted in a 1-inch acrylic centrifuge tube using open cell foam to hold the tissue in place. The tube was filled with normal saline to reduce the potential for projection over ranging. Scans were acquired at 130 kVp, 190 mA with a 0.125 mm Cu filter using 300 projections resulting with isotropic 25 μm isotropic resolution.

### CT registration and stereotaxic orientation

The DMBA stereotaxic space was defined using cranial landmarks extracted from a micro-CT dataset of the mouse skull. Using 3D Slicer, we manually placed ∼100 landmarks each along the sagittal suture, coronal suture, and lambdoid suture. A curve of best fit was calculated through each suture. For the sagittal suture, a polynomial curve was fit. Because of their more complex shape, the coronal and lambdoid sutures were fit with the discrete cosine transform ^45XX^. Before finding the curve intersections, we registered the CT scan to the DMBA MRH atlas volume in two stages. First, we created a brain tissue mask of the CT volume with thresholding and manual edits. This mask was affinely registered to a mask of the DMBA DWI volume. Then, we refined the registration using the Slicer Landmark Registration module with a Thin Plate Spline transform. With the CT scan aligned with the MR and in RAS orientation, we defined bregma as the intersection of the sagittal and coronal best fit curves, projected to the surface of the skull. We define lambda as the intersection of the sagittal and lambdoid best fit curves, projected to the surface of the skull. To fully define the stereotaxic space (to find the correct left-right rotation so the axial slices are not skewed), we placed a third landmark at the decussation of the anterior commissure, as identified by the RCCF label set that has been registered to the atlas volume. The volume was rotated such that bregma and lambda lay on the same axial slice, and all three landmarks lay on the same sagittal slice. The volume was then translated so that its origin is at bregma. The process was repeated on all five specimens. The average bregma-lambda distance was 4.31 ±0.31 mm. We choose the largest specimen 230328-4:1 with bregma-lambda of 4.67 mm as the reference for the atlas.

### Light Sheet Microscopy Acquisition

Upon completion of MR/DTI acquisition, the brains were removed from the skull, placed in buffered saline and shipped to LifeCanvas Technology (Cambridge, Mass.) for light sheet imaging. Formalin fixed samples were preserved using SHIELD following the outline described in ^53^. Each brain was incubated in 20mL of SHIELD-off solution for 4 d followed by 1 d of incubation in 20mL SHIELD-on solution. The meninges then were removed from each sample. Samples were incubated in Clearing Buffer A (LifeCanvas Technologies) overnight then actively delipidated using a LifeCanvas Technologies SmartClear II Pro device for 6 days using stochastic electrotransport ^54^. After depilation the samples were washed in PBS with 0.1% Tween20 for 1 d to remove SDS. For immunolabeling the samples were incubated in SmartLabel Primary Sample Buffer (LifeCanvas Technologies) overnight with an additional 5-6 h incubation with fresh buffer before primary immunolabeling in a SmartLabel device employing eFLASH technology which integrates stochastic electrotransport ^54^} and SWITCH ^55^ for 14 h. The samples were then washed in PBS for7-8h before overnight fixation in 4% paraformaldehyde followed by incubation in secondary labeling buffer at 37 C with two refreshes over the course of 7-8h before secondary labeling in the SmartLabel device.

**Table M1.**
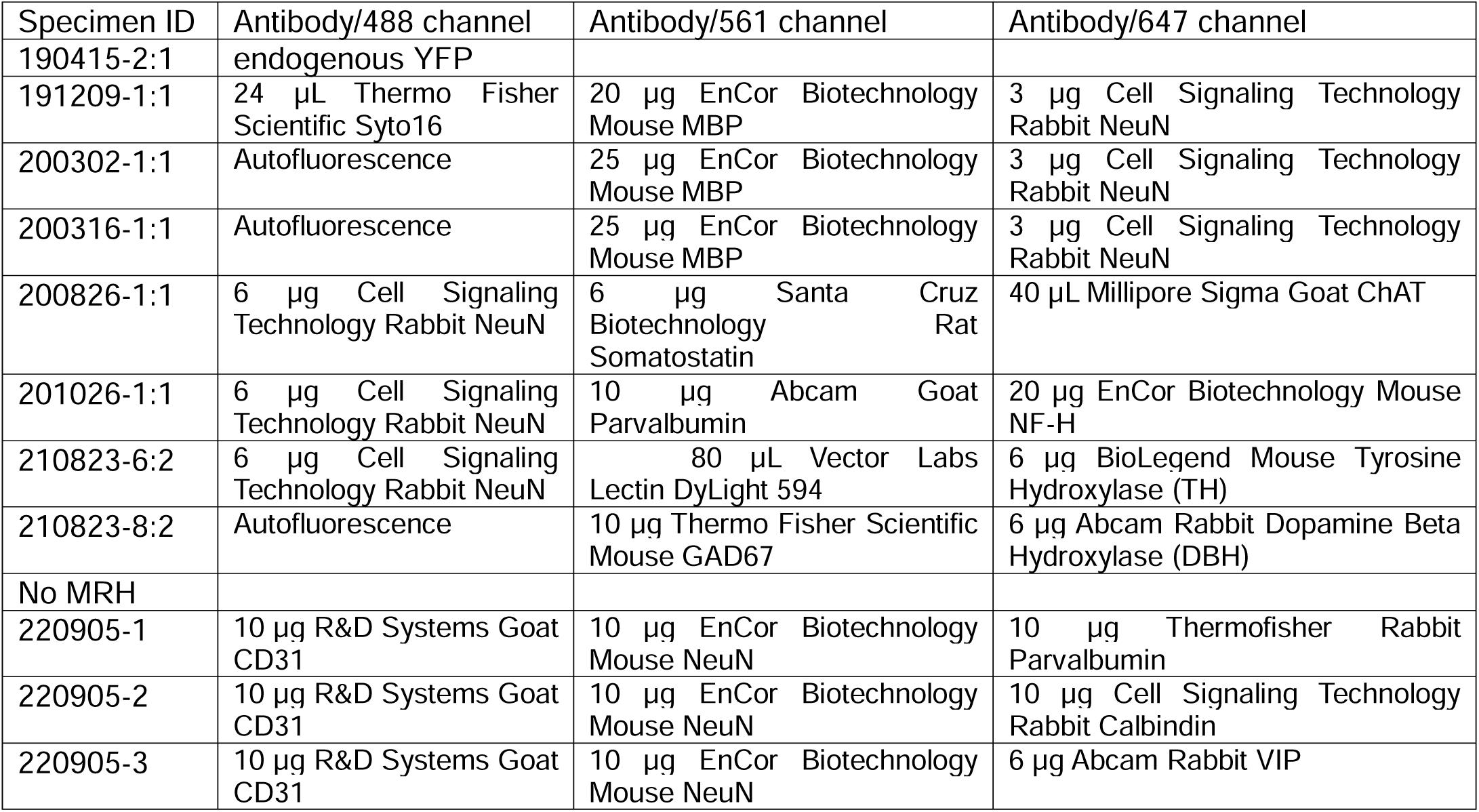
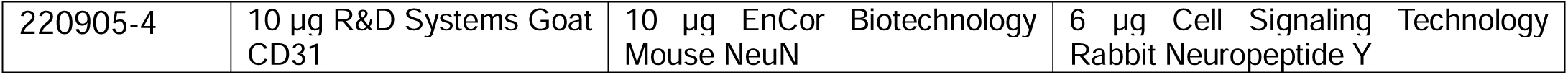
Antibodies used for light sheet microscopy.

### LSM Registration to DMBA

There are three registration pathways for LSM data, depending on the status of the specimen from which it came.

Light sheet volumes of specimens in the combined MRH w LSM template group (Table S2) were aligned to the MRH volume of the same specimen using methods outlined in detail in ^2^. This is a three-step registration process, starting with a coarse manual landmark registration in Slicer to account for the major tissue tearing, nonlinear swelling, and distortions. There are generally distortions throughout the brain, but they are most extreme in olfactory bulb and brainstem. DWI was the target volume contrast and NeuN was used as the moving volume contrast. Registration was performed at 15 μm resolution. The next step was a two-stage diffeomorphic registration using the antsRegistration from ANTs ^51^. Stage one used the BSplineSyN transform and Cross Correlation metric. This result was pushed to a refinement stage using the Symmetric Normalization (SyN) transform and Mattes metric. These transforms as well as those computed to create the DMBA template were then applied to the full resolution LSFM volume to warp into the DMBA stereotaxic space.

A second group of four LSM data sets (Table S2, specimens 220905-1:1-4:1) was acquired on specimens for which there were no MRH scans. The previous approach would not work since the first stage correction to an MRH volume was not possible. We instead created a group template using SAMBA as was used to create the DMBA MRI template. LSM NeuN volumes were down sampled to 45um. We performed N4 bias correction ^56^ to remove staining bias, and applied a log-scale to accommodate the dynamic range. Because we do not have an existing LSM atlas volume, one individual was arbitrarily chosen as the target for the rigid transform initialization stage. After all transforms were calculated, they were applied to the LSM volumes at 15 μm without the bias-correction or log-scaling. This LSM template was then registered to the MRH atlas using the pipeline described in the paragraph above, with this NeuN template as the moving volume and the DMBA DWI template as the registration target. Because both are average volumes, some individual variability is smoothed out and registration is more successful. This approach also only requires one manual landmark registration job instead of N, reducing the amount of human error in the results.

Finally, this NeuN template registered to DMBA provided a gateway to register arbitrary new LSM volumes into the common space. Volumes were down sampled to 45um, bias-corrected, and log-scaled. A rigid and affine initialization to the NeuN template were performed, followed by a diffeomorphic registration using the Cross Correlation metric and Symmetric Normalization (SyN) registration. This transform stack (rigid initialization, affine initialization, diffeomorphic registration to NeuN template) plus all transforms from the NeuN template to the DMBA DWI template were applied to the full-resolution version of the input volume to warp it into the DMBA stereotaxic space. While this approach does not have the same accuracy as the other two presented, it is an effective solution to warp arbitrary native-space light sheet volumes (that include NeuN or autofluorescence as a channel) into the DMBA stereotaxic space.

### Image Processing

Image processing was facilitated by the BigImage environment described in ^3^. 3D slicer (https://www.slicer.org/) and Imaris 10.0 and 10.1 (https://imaris.oxinst.com/) were used in preparing figures. A dedicated server receives LSM data via Globus and converts the three channels in each specimen (∼ 300GB/ channel) to tiff stacks. Raw (32 bit) images from the MRH pipelines described above and tiff stacks from the Nikon MicroCT and LifeCanvas were converted to ims format using the file converter (ImarisFileConverter 10.0) provided by Oxford Instruments. When image comparisons were made (e.g. Figure 1,2,3,6), all the volumes in a figure were loaded into Imaris at once. This was accomplished on one of our (4) high performance Dell servers each of which has 2TB of memory. Imaris is designed for 4D imaging with independent interactive adjustment of look up tables for each volume. The meta data on each volume of DMBA has been adjusted to place the original at bregma as described above. Imaris accommodates the varied spatial resolutions (LSM@ 1.8×1.8× 4.0 μm, TDI @ 5 μm, MRH @ 15 μm, microCT @ 25 μm) by scaling the data internally. Imaris provides a fiducial bar that was captured in each image being compared and carried through in assembling figure to provide provenance on spatial resolution. When assembling figures with multiple resolution, care was taken to load the highest resolution data first which assures that slices being compared are done so at their native slice thickness unless purposely overridden. Minimum, maximum and gamma are adjusted to map the histogram of the volume to the available gray (or color) scale. Individual frames are captured at at least the Nyquist resolution of the image being shared. Final figure assembly takes place in Photoshop and Illustrator were gray or color scale from the individual components of the images (from different modalities) are fine-tuned by adjusting the minimum, maximum and gamma to the available range in the figure. In some cases, scale bars in the final composite are then edited out.

## Acknowledgements

This work was supported by grant R01NS120954-01A1 from the National Institute of Neurological Disorders and Stroke (to FCY and GAJ) and grant R01AG070913-01 from the National Institute on Aging (to RWW and GAJ). We are grateful to Justin Gladman in the Duke Shared Material Instrumentation Facility for crucial assistance in obtaining the microCT data. We are grateful to Tatiana Johnson for editorial and formatting assistance.

## Author contributions

All authors reviewed the manuscript. HM did the majority of the image processing including creating the QSDR volumes, labeling, analysis of CT images, mapping CT to QSDR volume, definition of bregma and lambda. RA assisted HM and YT in image processing, performed quality assurance checks on MRH to LSM registration. JCC created the pipelines for image acquisition and reconstruction and denoising. He maintains the big data infrastructure used in image processing. KJH performed statistical analysis and generated figures. YQ performed tissue perfusion,staining and preparation for scanning. She maintains the tissue data archive. YT developed the pipelines for light sheet to MRH registration and neuron counting. She performed data analysis and prepared figures. RWW assited in project definition, provided advice on neuroanatomy, stereology and sterotaxic framework, and assisted in preparing the manuscript. He provided funding through R01AG070913-01 (to RWW and GAJ). FCY is the creator of DSI Studio. He assisted in image processing and preparing figures. He advised in creation of the image processing pipeline and provided funding through R01NS120954-01A1. LEW provided expertise in neuroanatomy and stereotaxic frameworks. He provided major assistance in data analysis, manuscript preparation and project direction and shares the position of senior author with GAJ. GAJ conceived and directed the project, wrote the manuscript and provided funding in collaboration with RWW and FCY.

## Correspondence

All correspondence and requests for additional data should be addressed to G Allan Johnson.

## Open Access

This work is licensed under the creative common CC BY-NC-SA. This license enables users to distribute, remix, adapt, and build upon the material in any medium or format for noncommercial purposes only, and only so long as attribution is given to the creator. If you remix, adapt, or build upon the material, you must license the modified material under identical terms. CC BY-NC-SA includes the following elements: BY: credit must be given to the creator. NC: Only noncommercial uses of the work are permitted. SA: Adaptations must be shared under the same terms.

