## Supplemental Figures and Tables for "An Open Resource: MR and light sheet microscopy stereotaxic atlas of the mouse brain"

**Supplement**

**Table S1.** Representative mouse brain atlases with elements similar to this work.

| Ref | Strain/N | In/Ex  vivo | in/  out | Stereo | Voxel  dimension (μm) | Contrast/Resolution Index | Field  Tesla | Labels |
| --- | --- | --- | --- | --- | --- | --- | --- | --- |
| ^1^ | B6 | Ex | in |  | 39x39x156 | T2*  DTI/25,287 | 9.4 | NA |
| ^2^ | B6/NA | Ex | out |  | 60x60x60  histology | T2  MBP,GFAP,Nissl,AChE | 11.7 | NA^1^ |
| ^3^ | 129S1/  Svimj | Ex | out | yes | 60x60x60 | T2 | 7 | 42 |
| ^4^ | B6/10 | Ex | out |  | 47x47x47 | T2* | 17.6 | 20 |
| ^5^ | B6,CD1  129S1 | Ex | out | yes | 54x54x54  38x38x38 | T2  CT | 7 | NA^2^ |
| ^6^ | B6 | Ex | in |  | 21.5x21.5x21.5  43x43x43 | T1  T2 | 9.4 | 34 |
| ^7^ | B6/40 | Ex | in |  | 32x32x32 | T2 | 7 | 62 |
| ^8^ | B6/  BXD/11 | Ex | in |  | 43x43x43 | T1,T2 | 9.4 | 34 |
| ^9^ | B6/9 | In  Ex | in | yes | 50x50x125  62.5x62.5x62.5 | T2  T2,DTI/24,576 | 9.4  11.7 | 46 |
| ^10^ | B6/12 | Ex | in |  | 21.5x21.5x21.5  43x43x43 | T1,T2*  T2 | 9.4 | 37 |
| ^11^ | B6/8 | Ex | in |  | 43x43x43 | T1,T2  DTI/75,465 | 9.4 | 37 |
| ^12^ | B6/18 | Ex | out | yes | 30x30x30 | T2* | 16.4 | 187^3^ |
| ^13^ | B6/18 | Ex | out |  | 100x100x100 | DTI/30,000 | 16.4 | 187^3^ |
| ^14^ | B6 | Ex | in |  | 43x43x43 | DTI/1.5x10^6^ | 9.4 | 296 |
| ^15^ | B6/23 | Ex | out | yes | 60x50x200 | T2* | 7 | P_F |
| ^16^ | B6/12 | Ex | out |  | 33x33x33  8.6x100x8.6  8.6x33x8.6 | Flash  DAPI | 9.4 | ABA |
| ^17^ | RjOrl:  SWISS | In | In |  | 120x120120 | T2 | 7 | ABA |
| ^18^ | B6/6 | In | in |  | 120x120x400 | SANDI 8 shell 40 direct |  | ABA |
| ^19^ | B6/12 | In  Ex | in |  | 150x150x400-500  100x100x500 | T2/MT/OGSE  DTI/3076 | 9.4 | NA |
| ^20^ | B6/12 | Ex | in | yes | 78x78x78  125x125x125  22.6x22.6x22.6  4.8x4.8x3.8 (20) | T2  DTI/30,720  uCT  AutoF |  | ABA |
| ^21^ | B6/30 | In  Ex | in |  | 100x100x100  100x100x100 | DTI/2000  DTI/5,000 | 7  11.7 | ABA |

^1^ The total number of animals studied was not specified.

^2^ Several different strains were scanned. The total number of each was not specified.

^3^ In a series of four papers, the basal ganglia, cerebellum, cortex, and hippocampus were delineated with 35,38,74,40 delineations (total =187) on the same data set.

**Table S2.** Specimens used to construct the atlas.

|  | **Specimen ID** | **Body Wt** |
| --- | --- | --- |
| **MRH w LSM** | 200302-1:1 | 30.2 |
|  | 200826-1:1 | 24.6 |
|  | 201026-1:1 | 24.2 |
|  | 210823-8:2 | 26.2 |
|  | 210823-6:2 | 25.9 |
|  | 190415-2:1 | 23.6 |
| **MicroCT** | 230328-4:1 | 30.4 |
|  | 201202-1_1 | 27.1 |
|  | 201202-1_2 | 26.8 |
|  | 201202-1_3 | 27.5 |
|  | 201202-1_4 | 26.3 |
| **LSM** | 220905-1:1 | 26.8 |
|  | 220905-2:1 | 29.7 |
|  | 220905-3:1 | 28.4 |
|  | 220905-4:1 | 24.4 |

**Table S3.** MRH volumes of the average atlas included with the release, the algorithms used to generate them and the size of each data volume.

| **Average MRH Library** |  |  |
| --- | --- | --- |
| File Name | Algorithm | Volume (GB) |
| DMBA_MGRE_AVG_NOMASK | Average Echo0-3-no Mask | 4.2 |
| DMBA_MGRE_AVG | Average Echo0-3 | 4.2 |
| DMBA_MGRE_M0 | TE=5.4 ms | 4.2 |
| DMBA_MGRE_M1 | TE=10.9 ms | 4.2 |
| DMBA_MGRE_M2 | TE=16.3 ms | 4.2 |
| DMBA_MGRE_M3 | TE=21.7 ms | 4.2 |
| DMBA_DWI | Average all b_N_ | 4.2 |
| DMBA_AD | DTI | 4.2 |
| DMBA_FA | DTI | 4.2 |
| DMBA_MD | DTI | 4.2 |
| DMBA_RD | DTI | 4.2 |
| DMBA_ISO | GQI | 4.2 |
| DMBA_QA | GQI | 4.2 |
| DMBA_CLR_GQI | GQI | 1.6 |
| DMBA_TDI_3X | TDI ^22^ | 42.8 |
| DMBA-MicroCT | CT | 3.65 |

Abbreviations are defined in main text.

**Table S4.** Light sheet volumes included in the DMBA and immunohistochemistry labels used in each channel of the SPIM.

| **LSM-MRH** | **Channel** |  |  |
| --- | --- | --- | --- |
| Specimen ID | 488 nm | 561 nm | 642 nm |
| 190415-2:1 | End YFP | AutoF |  |
| 191209-1:1 | Syto | MBP | NeuN |
| 200302-1:1 | AutoF | MBP | NeuN |
| 200316-1:1 | AutoF | MBP | NeuN |
| 200826-1:1 | NeuN | SST | ChAT |
| 201026-1:1 | NeuN | Parvalbumin | NFH |
| 210823-6:2 | NeuN | Lectin | TH |
| 210823-8:2 | AutoF | DBH | GAD67 |
| **LSM-No MRH** |  |  |  |
| 220905-1 | CD31 | NeuN | PV |
| 220905-2 | CD31 | NeuN | Calbindin |
| 220905-3 | CD31 | NeuN | VIP |
| 220905-4 | CD31 | NeuN | NPY |

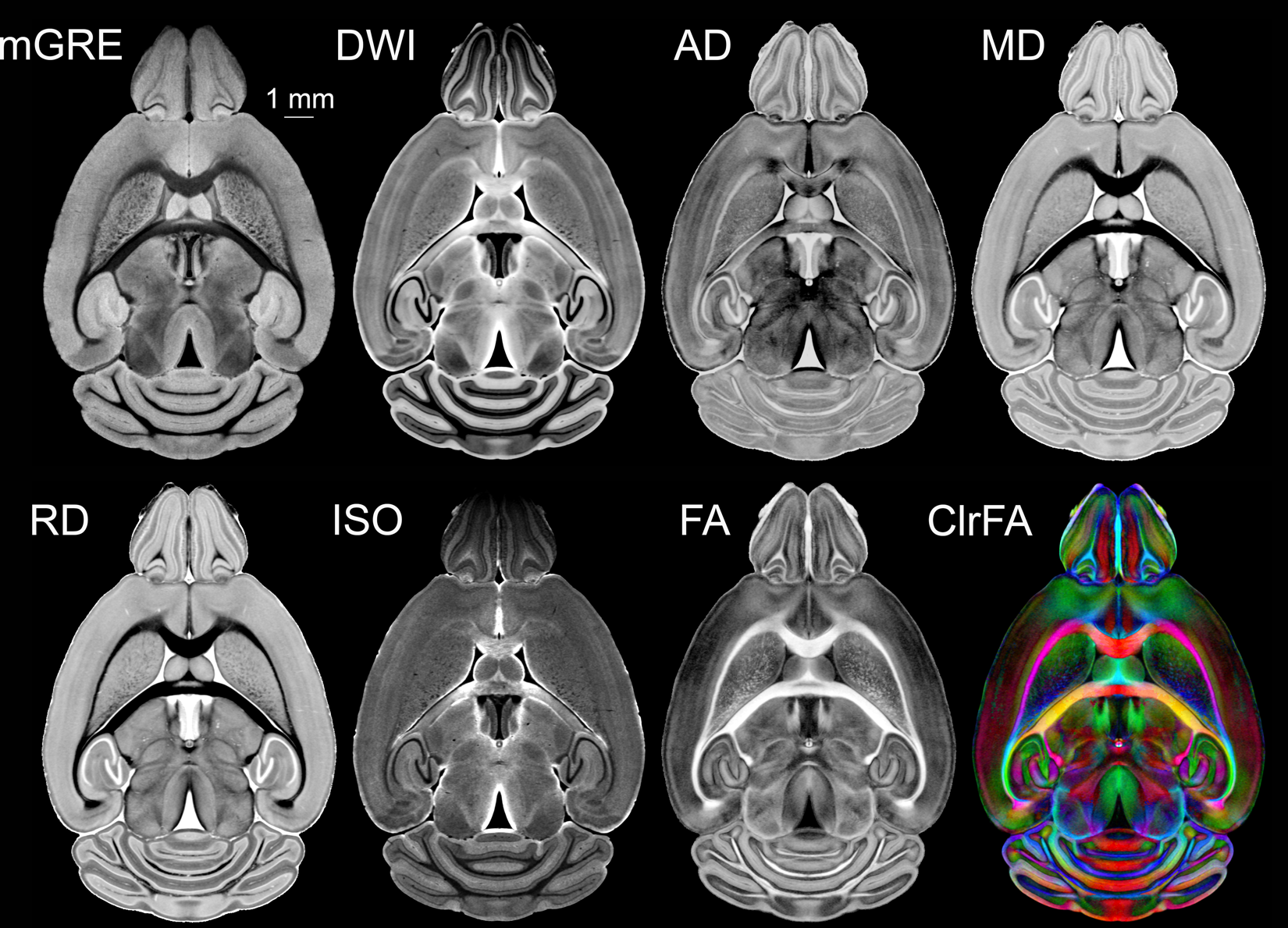

**Figure S1. Multiple MRH volumes derived from the mGRE and dMRI volumes highlight different histological features.** The same plane from the mGRE volume and selected images derived from the dMRI volumes demonstrate the wide range of contrast delineating differing brain anatomy. Abbreviations are those used in the main text.

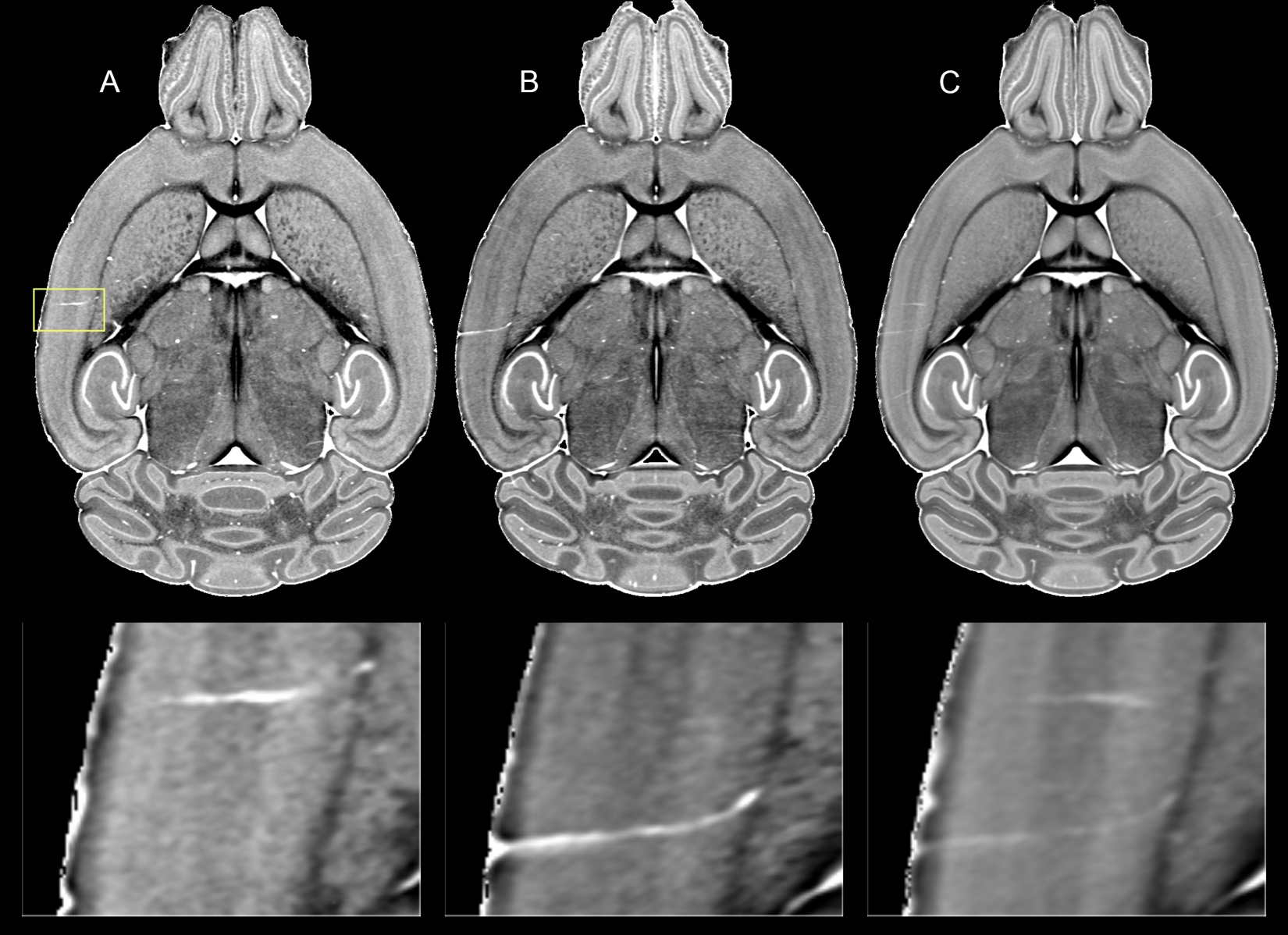

**Figure S2. Creation of the minimum deformation averages produces vascular artifacts.** A) MD image from specimen 200302-1:1 showing penetrating vessel (yellow box, magnified in bottom row). B) MD image from the same level in specimen 200826-1:1 showing a second penetrating vessel in a similar (but not identical) position. C) MD image at the same level in the MDT average atlas showing superposition of both vessels.

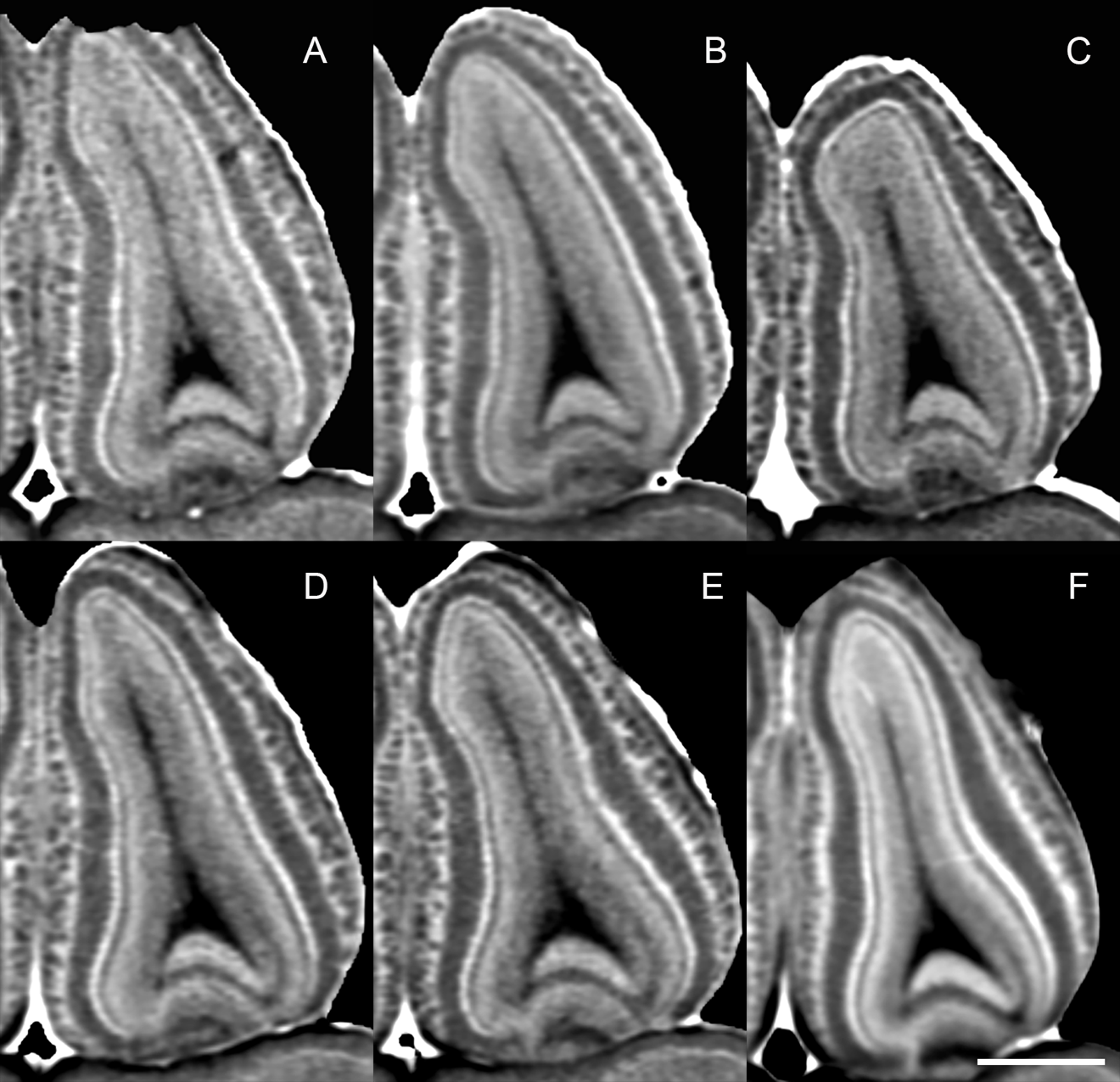

**Figure S3. Cellular layers of the olfactory bulb are enhanced in the MDT average while the glomeruli are blurred.** Magnified section of the olfactory bulb in the MD images from the individual specimens: A) 200302-1:1; B) 200826-1:1; C) 201026-1:1; D) 210823-8:2; E) 210823-6:2; and	F) the MD average. The scale bar is 0.8 mm. Note the olfactory glomeruli are resolved in the individual images but blurred in the average, but the cellular layers remain distinct and are enhanced.

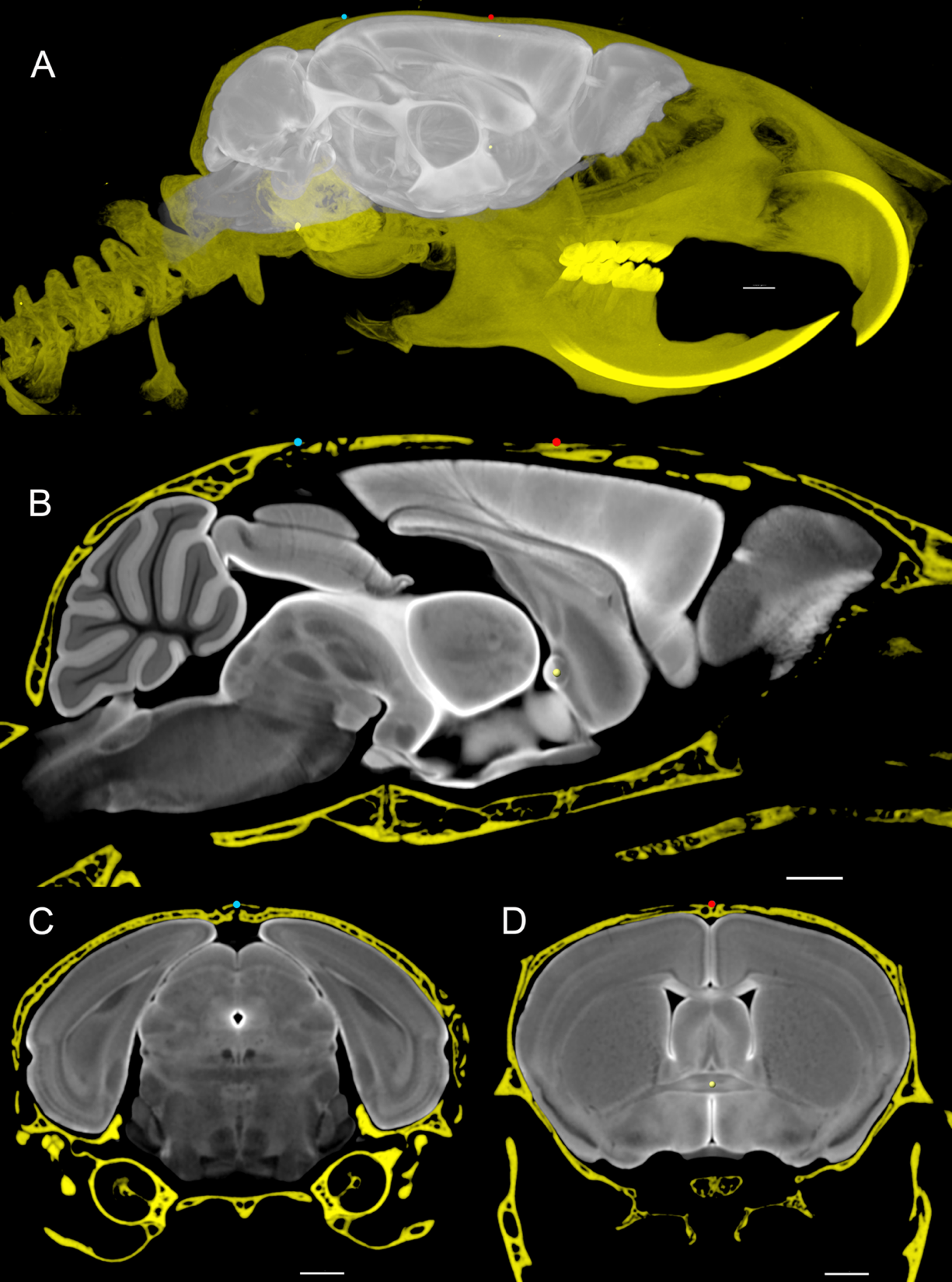

**Figure S4. Bregma and lambda are defined by a 3D micro-CT image aligned to the MDT MRH volume.** A) The entire CT volume is shown to appreciate the location of the incisor. Bregma (red dot) and lambda (blue dot) are defined on the CT volume which has been registered to the space defined by the MRH volumes acquired with the brain in the skull-DWI in this example. B) Sagittal slice through the registered stereotaxic space displayed at higher magnification. The cardinal coronal plane is defined by a plane intersecting bregma and the midpoint of the anterior commissure (yellow dot). C) Coronal slice @ lambda. D) Coronal slice @ bregma. Scale bars are 1 mm.

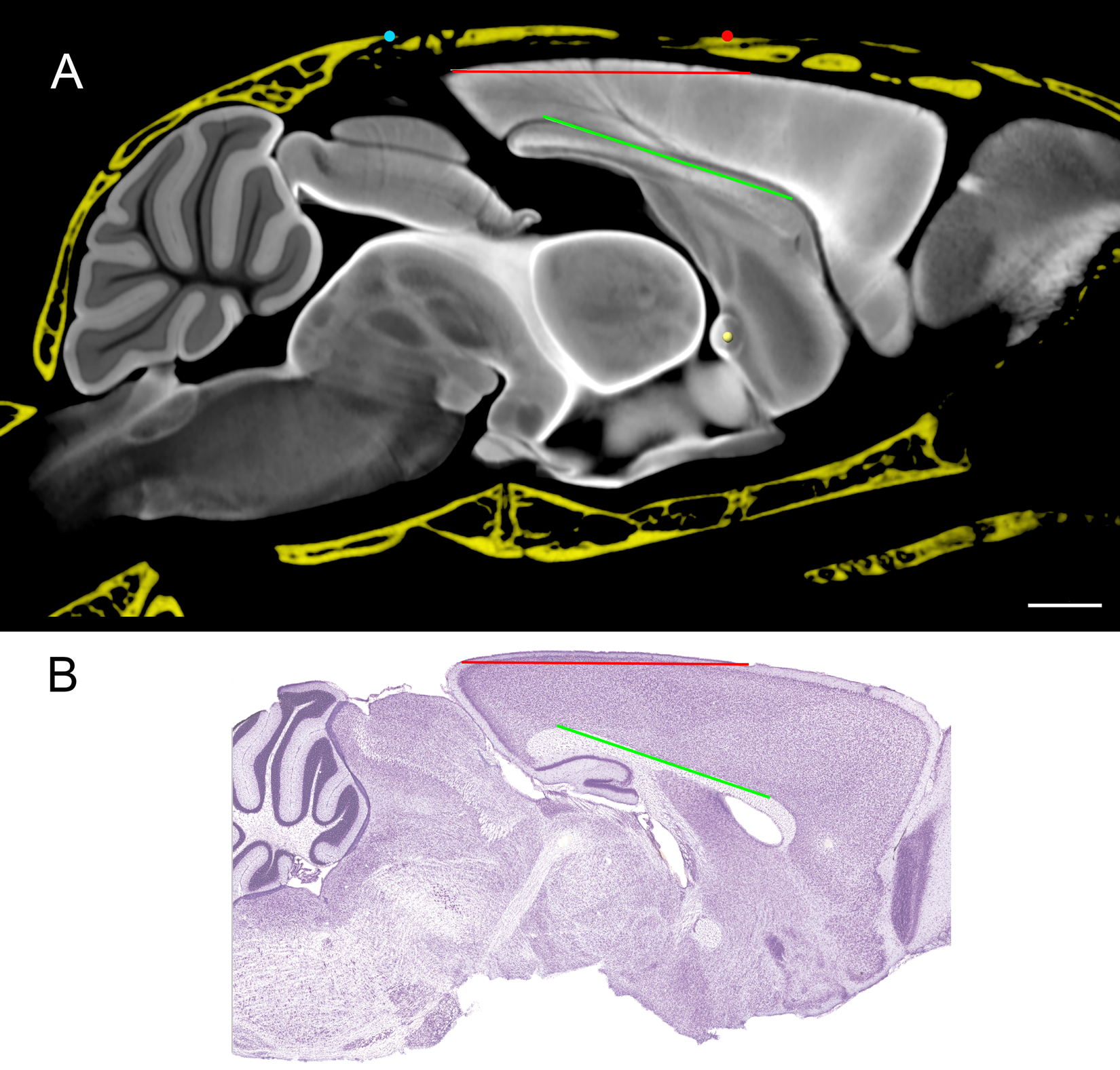

**Figure S5. DMBA is aligned to the Franklin Paxinos (FP) atlas [ref. 23]**. A) Sagittal DWI from DMBA in which the flat skull angle (red) is -0.16 degrees. The angle of the corpus callosum (green) is -18.4 degrees. B) Sagittal Nissl section (Plate 104) from FP ^23^. The flat skull angle (red) is -0.49 degrees. The angle of the corpus callosum (green) is -18.7 degrees. Scale bar is 1 mm. The images have been scaled so their horizontal axes are the same to allow visual comparison.

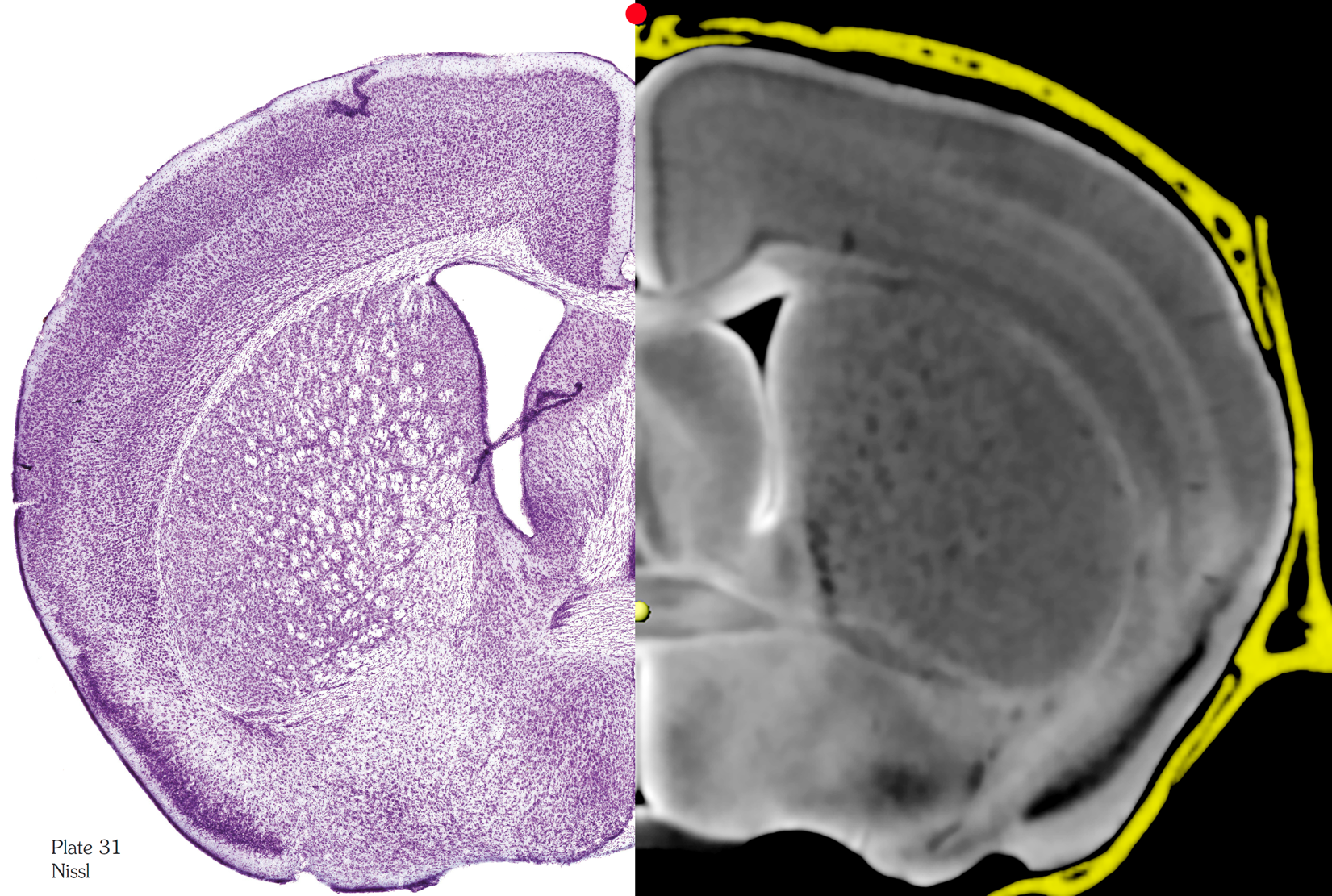

**Figure S6. Coronal slices at bregma are in good agreement between the Franklin Paxinos (FP) atlas [ref. 23] and the DMBA**. Left, Nissl (plate 31) from the FP atlas^23^ at Bregma and DWI at Bregma from DMBA confirming that the orientation of the two atlases is very similar. The two images have been scaled so the vertical dimension are the same to allow visual comparison of anatomic landmarks. The height/width is different in the two since the DMBA is in the skull.

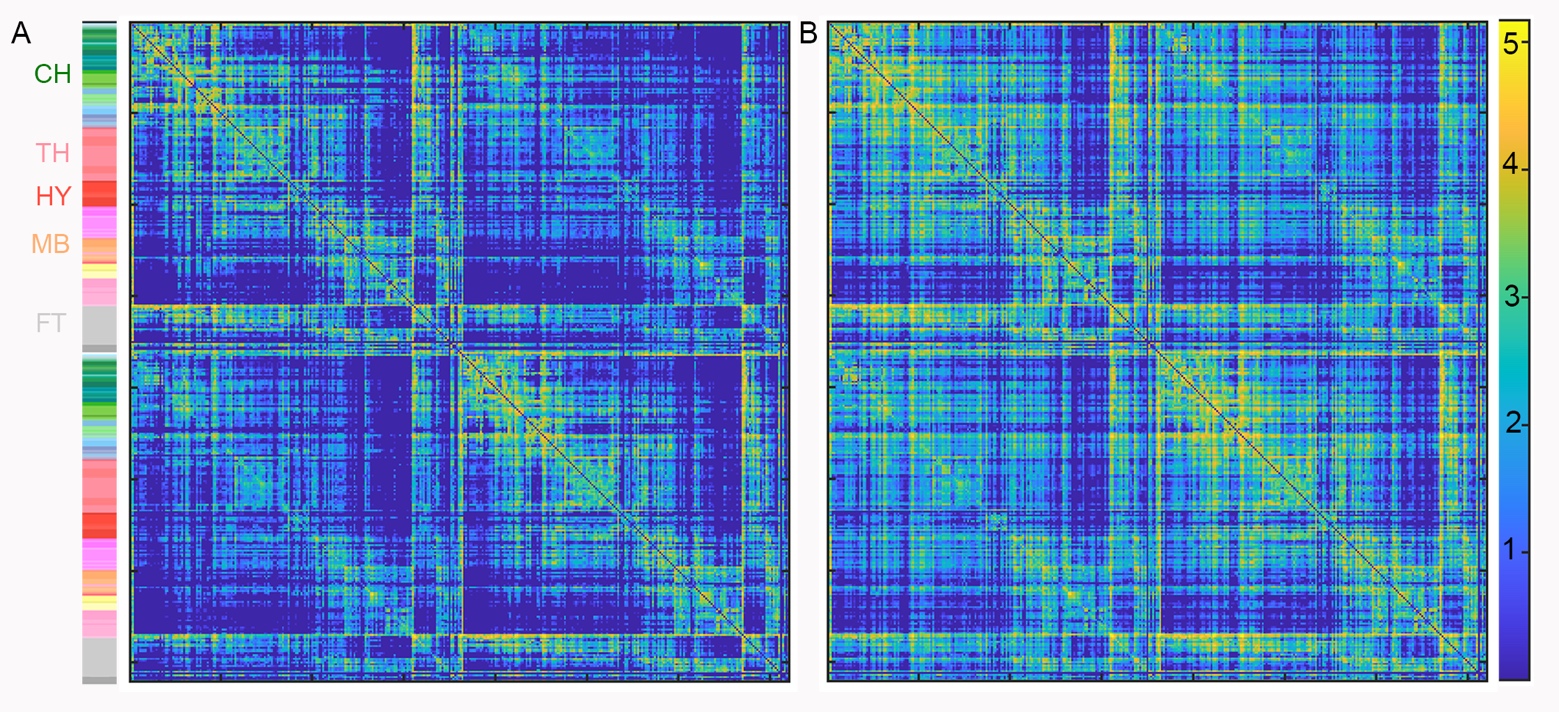

**Figure S7. A) Connectome of a single C57BL/6J 90-day male mouse and B) QSDR of five C57BL/6J mice show connection strength within nodes in left hemisphere (upper left quadrant of each matrix), within the right hemisphere (lower right quadrant of each matrix), and between hemispheres (upper right and lower left quadrant of each matrix).** The color bar on the left localizes the ROIs from Cerebrum (CH), Thalamus (TH), Hypothalamus (HY), Midbrain (MB), and Fiber tracts (FT) on the left hemisphere following the color standards employed by the Allen Brain Atlas. The strength of connection between these nodes is indicated by the color bar on the right which is a Log_10_ scale covering 4-5 orders of magnitude. The average of five specimens in B highlights many more subtle connections than are visible in the connectome of the single specimen.

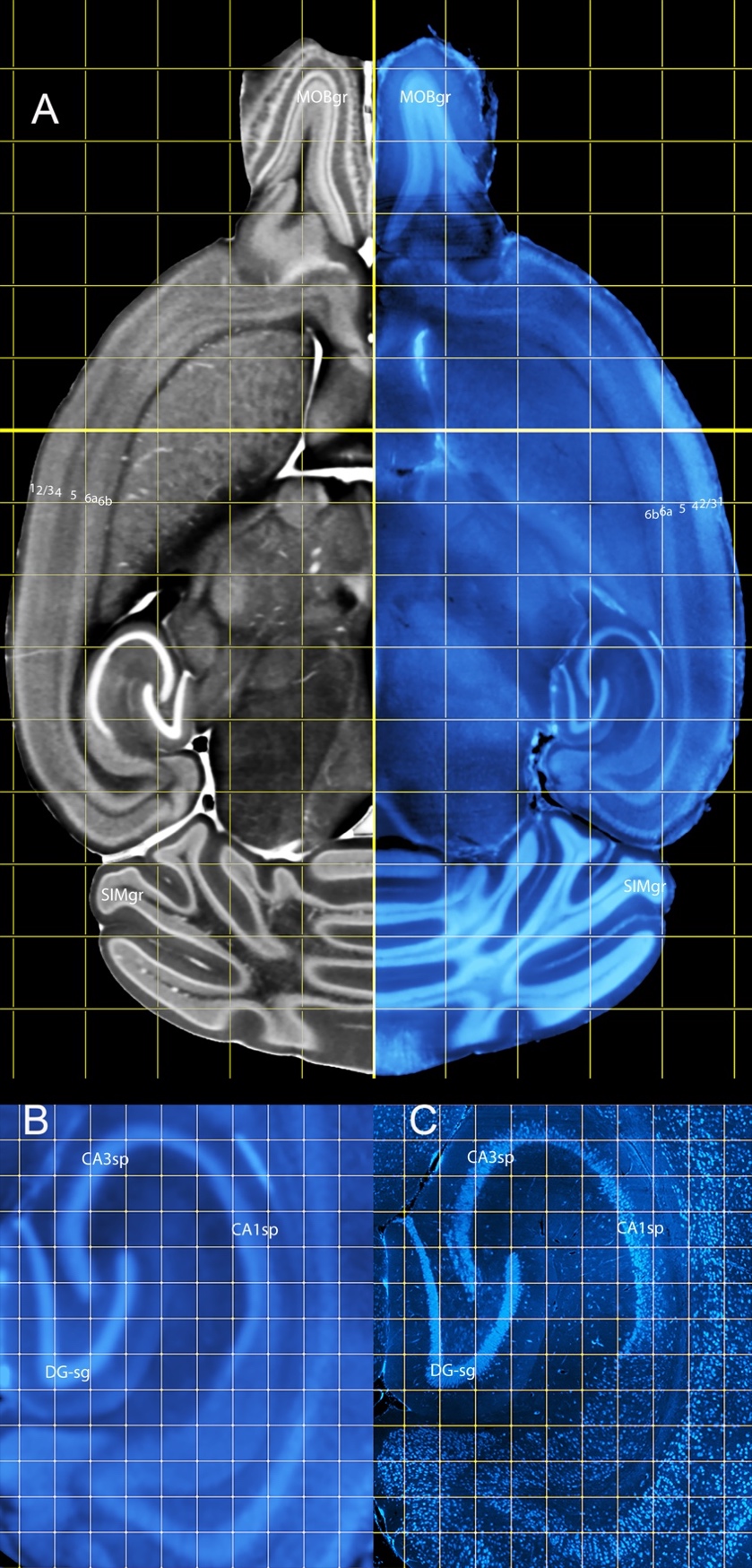

**Figure S8. Minimum deformation Template (MDT) NeuN (A-Right) registered to DMBA_MD (A-Left) links MRH and LSM libraries.** The NeuN-MDT provides a gateway for light sheet users to register their data into stereotaxic space. CA1sp, field CA1 pyramidal layer; CA3sp, field CA3 pyramidal layer; DG-sg, dentate gyrus, granule cell layer; MOBgr, main olfactory bulb, granule layer; SIMgr, simple lobule granular layer; numbers indicate cortical layers. Grid array in A is 1 mm spacing and 200 μm spacing in B and C.

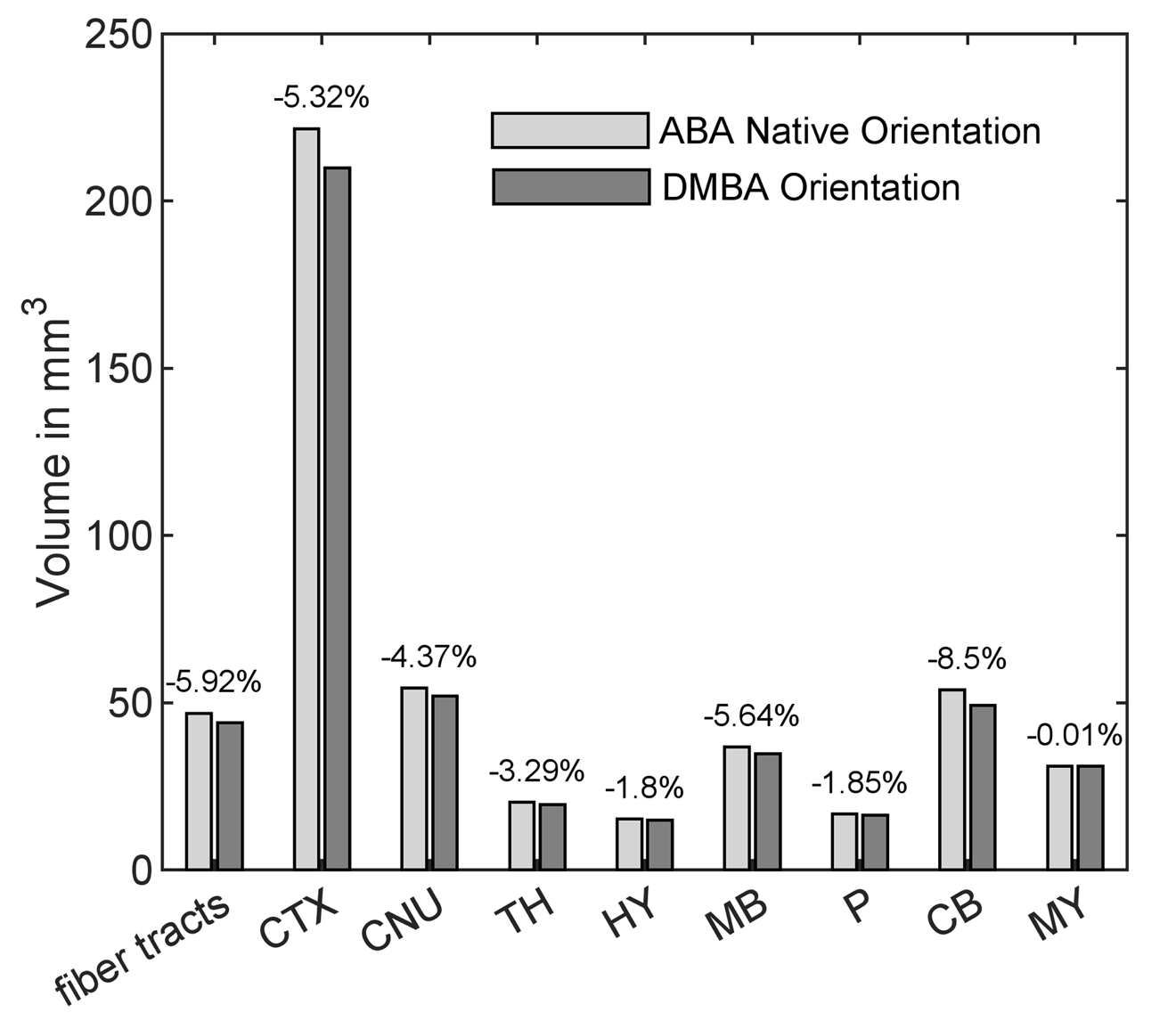

**Figure S9. Differences in volume for some of the larger anatomic structures when the ABA ROI are transformed into the stereotaxic space demonstrated in Figure 5.** For many of the larger structures, the differential swelling in X and Y results in overestimate of the volumes in ABA. Percentages are relative volumes differences between orientations. Note the variable impacts of distortion correction on different brain regions.

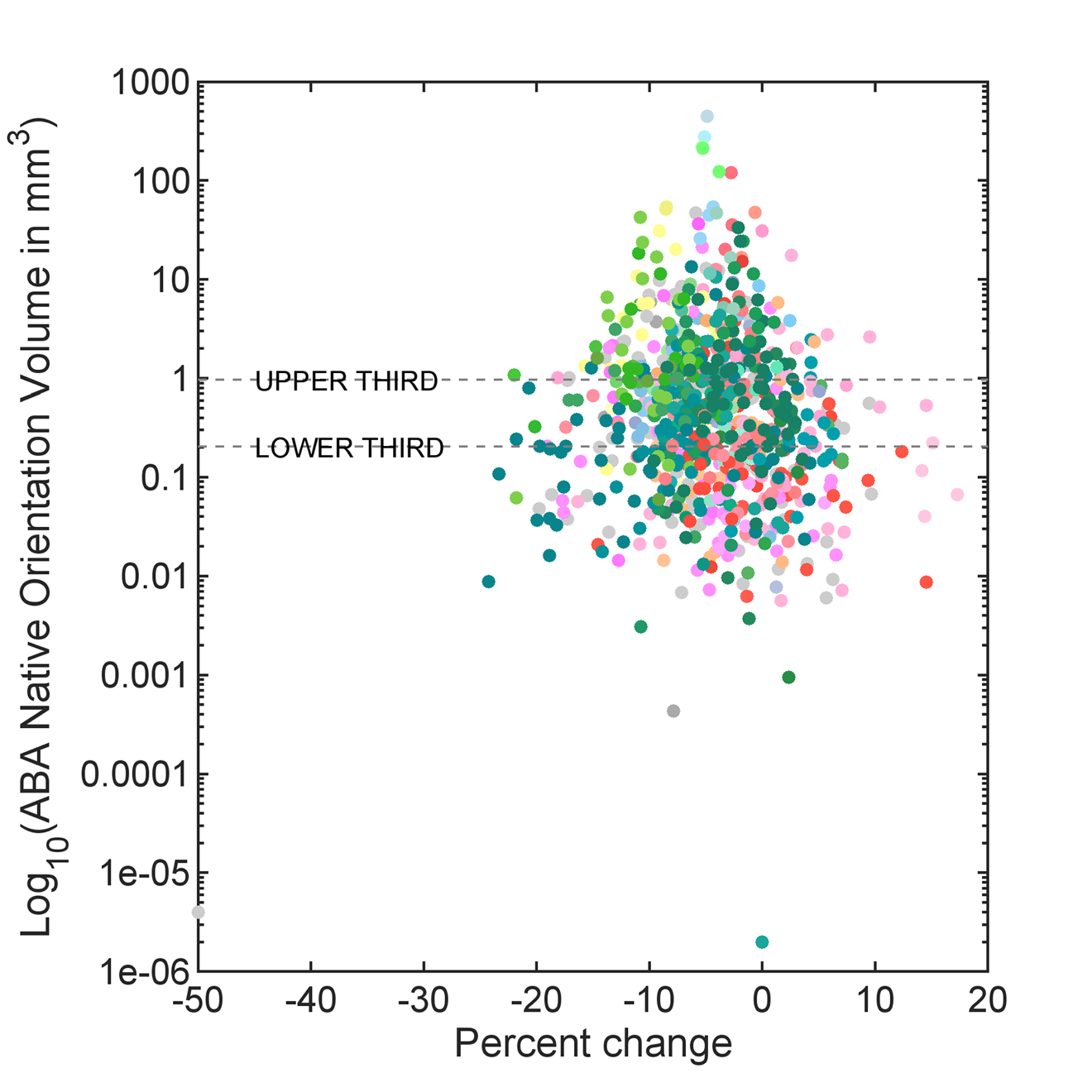

**Figure S10. Transform of ABA to stereotaxic space demonstrates that volume estimates are off by factors ranging between -25% to +15%.** Most of the ROI are larger in ABA consistent with non-uniform swelling following removing the brain from the skull. Colors are those used in the ABA labels.

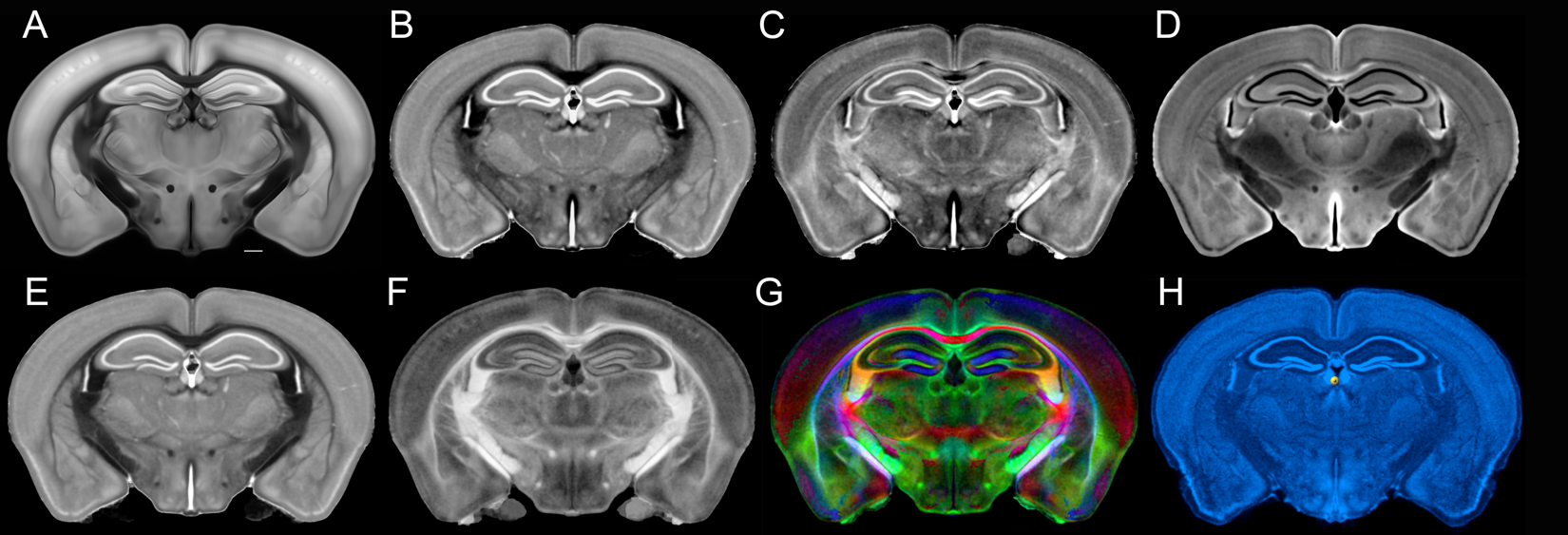

**Figure S11.** **The DMBA MRH atlas provides expanded structural detail not present in the ABA**. Coronal sections from A) The corrected ABA auto fluorescent image, B) MD, C) AD, D) DWI, E) RD, F) FA, G) GQI ClrFA, and H) corrected ABA Nissl section. Scale bar in A is 0.5 mm.

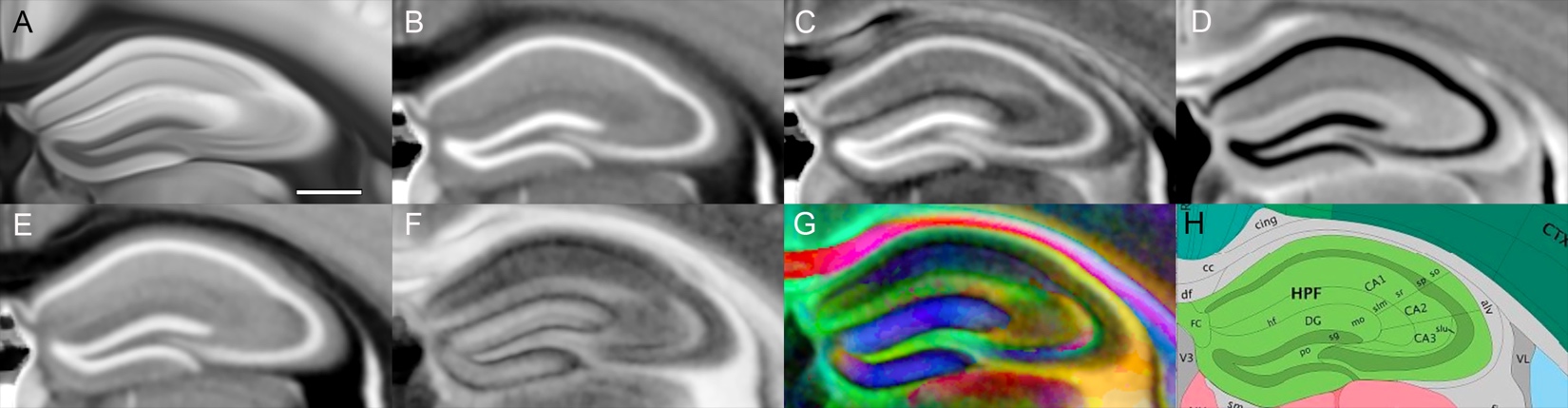

**Figure S12. Magnified view of hippocampus in a coronal plane demonstrates multiple, different cytoarchitectural features evident in the DMBA atlas.** A) The corrected ABA auto fluorescent image, B) MD, C) AD, D) DWI, E) RD, F) FA, G) GQI ClrFA, H) ABA labels. Scale bar in A is 0.5 mm.

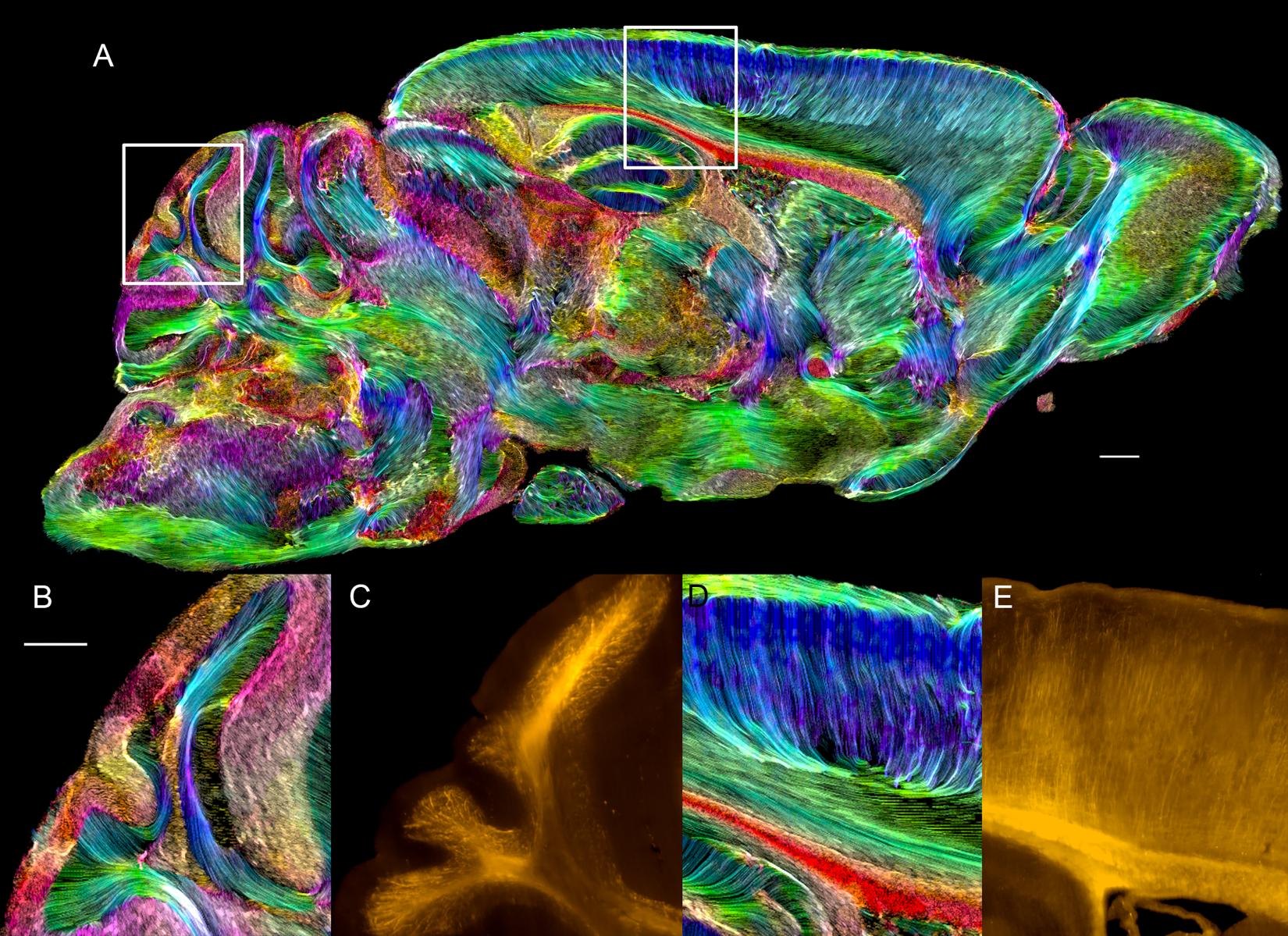

**Figure S13. Track density images resolve neurites, fascicles, and fiber tracts while providing spatial orientation.** DMBA_TDI is a super-resolution track density image at 5 μm isotropic resolution with color encoding indicating the local fiber orientation. A) The sagittal image @ 1.130 mm off the midline includes an area of cerebellum (left box) and neocortex (right box) which have been magnified and paired with a light sheet image (also part of the atlas) stained for myelin basic protein (Specimen 200316-1). Fascicles, fiber tracts, and colinear segments of neurites are prominent throughout the brain. Notably, TDI signal is not limited to cerebral white matter; panels **B** and **D** clearly show prominent fascicles in the subcortical white matter of the cerebellum and neocortex, respectively. These same regions stain intensely for myelin-basic protein, as shown in the registered LSM panels (**C** and **E**). In addition, there is robust TDI signal in the molecular layer of the cerebellar cortex, where the warm hues indicate the presence of unmyelinated parallel fibers of cerebellar granule cells coursing across the outer layer of cerebellar folia (see **B**). In neocortex, radially oriented TDI profiles indicate the co-linear arrangement of incoming thalamocortical afferents and cortico-cortical connections, which are also evident in the myelin-basic protein stain (cf. panels **D** and **E**). In **D**, it is likely the apical dendrites of cortical pyramidal neurons are also contributing to the strong radial signal apparent in TDI. The scale bar in A is 0.5 mm

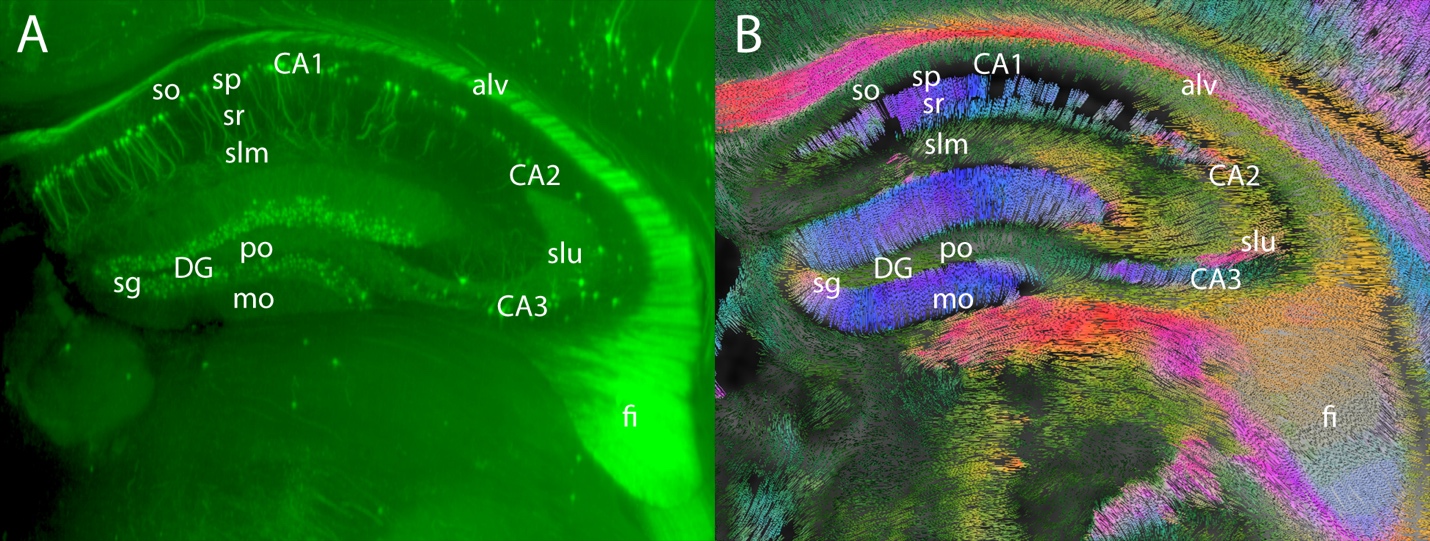

**Figure S14.** **Comparison of the hippocampus viewed in light sheet microscopy and track density imaging**. A) Light sheet image from Thy1/YFP (Specimen 190415-2_1 and B) tractography image from DMBA at the same level viewed in the coronal plane. The prominent TDI signal in B) colored purplish blue (neurites mainly in the dorsal-ventral axis) is likely associated with the dendrites of granule cells in the dorsal and ventral “blades” of the dentate gyrus, and the proximal apical dendrites of pyramidal neurons in fields CA1 and CA3 (stratum radiatum). Meanwhile, the hippocampal sublaminae containing the more distal apical dendrites (stratum lacunosum-moleculare) are more heterogenous with axonal projections and dendritic arborizations more broadly distributed in spatial orientation. Abbreviations: alv, alveus; CA1-3, Ammon’s horn fields 1-3; DG, dentate gyrus; mo, dentate gyrus molecular layer; po, dentate gyrus polymorph layer; sg, dentate gyrus granule cell layer; fi, fimbria; slu, stratum lucidum; slm, stratum lacunosum-moleculare; so, stratum oriens; sp, pyramidal layer; sr, stratum radiatum.

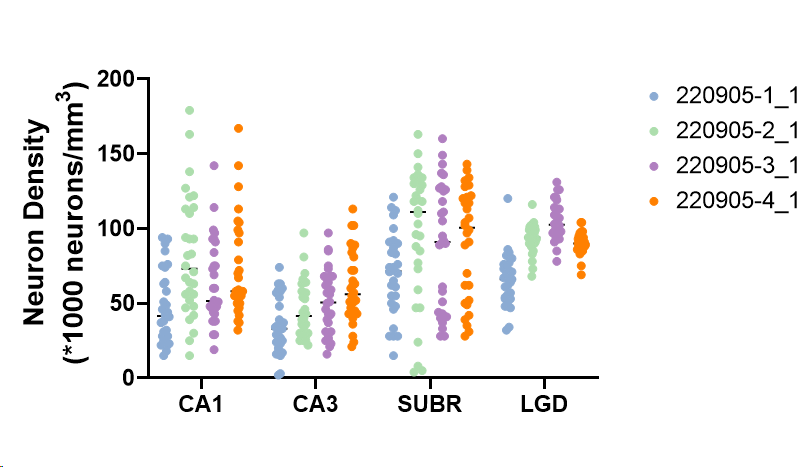

**Figure S15. Neuron density comparison across 4 male C57BL/6J mice.** Each point is the density of neurons in a 100 μm sampling cube obtained from NeuN-labeled specimens and imaged by light sheet microscopy (LSM). The cubes are randomly cast across the structure in the LSM volume after it has been geometrically registered to the specimen’s MRH image. Thus, the spread for each animal is an indication of the heterogeneity in neuron density within the structure. Abbreviations: CA1 and CA3, hippocampus Ammon’s horn fields 1 and 3, respectively; SUBR, subicular region; LGD, dorsal part of the lateral geniculate complex.

23.	Franklin KaP, Paxinos, G. The mouse brain in stereotaxic coordinates-third edition. San Diego: Academic Press; 2007.
